# Rare genetic variants in the IIS/mTOR signalling pathway identified in exceptionally long-lived individuals show shared in vitro effects associated with lifespan across species

**DOI:** 10.64898/2026.05.28.728260

**Authors:** Mara Neuerburg, Larissa Smulders, Erik B. van den Akker, Daniel Kolbe, Filippo Artoni, Isabell Brusius, Helena Hinterding, Laura Beltrame, Ramona Pahl, H. Eka D. Suchiman, Antonios Papadakis, Andreas Beyer, Marian Beekman, Almut Nebel, P. Eline Slagboom, Maarouf Baghdadi, Joris Deelen

## Abstract

**Background:** The increase in human lifespan without a proportional increase in healthspan imposes a substantial burden on individuals and society. Exceptionally long-lived individuals and members of long-lived families exhibit compression of multi-morbidity. Genetics, and in particular rare protein-altering variants, appear to play an important role in their longevity.

**Methods:** In this study, we employed a targeted pathway approach to provide functional evidence of the significance of rare variants in the insulin/insulin-like growth factor 1 signalling – mechanistic target of rapamycin (IIS/mTOR) signalling pathway identified in long-lived individuals. To this end, we used CRISPR/Cas9 to introduce these rare genetic variants into mouse embryonic stem cells (mESCs). We subsequently assessed several functional readouts that have previously been associated with lifespan regulation in model organisms and/or IIS/mTOR and mitogen-activated protein kinase/extracellular signal-regulated kinase (MAPK/ERK) signalling pathway activity.

**Results:** Functional characterisation revealed that the variants exhibit both shared and distinct effects on the signalling pathways. Principal component analysis of omics-based datasets showed that the variants clustered into two groups, a distribution that corresponds with the grouping observed for a subset of functional readouts. All variant mESC lines exhibited a downregulation in IIS/mTOR and MAPK/ERK signalling pathway activity as well as an increase in *Foxo3* expression and FOXO3 binding activity. We identified alterations in lipid and mitochondrial metabolism, including a reduction in mitochondrial DNA levels, which were mostly shared among all variants. All variant mESC lines exhibited a signature implying increased pluripotency. The effects on stress resistance and growth rate diverged between the two variant groups, with partially opposing effects. Group 1 demonstrated a reduced growth rate and increased resistance to a subset of stressors, while Group 2 demonstrated an increased growth rate and reduced resistance to a subset of stressors.

**Conclusions:** Here, we provide evidence that rare genetic variants in the IIS/mTOR and MAPK/ERK signalling pathways identified in long-lived human individuals result in shared functional effects associated with longevity in model organisms. These insights can serve as a foundation to better understand the role of rare variants in the insulin signalling network in the regulation of human longevity.

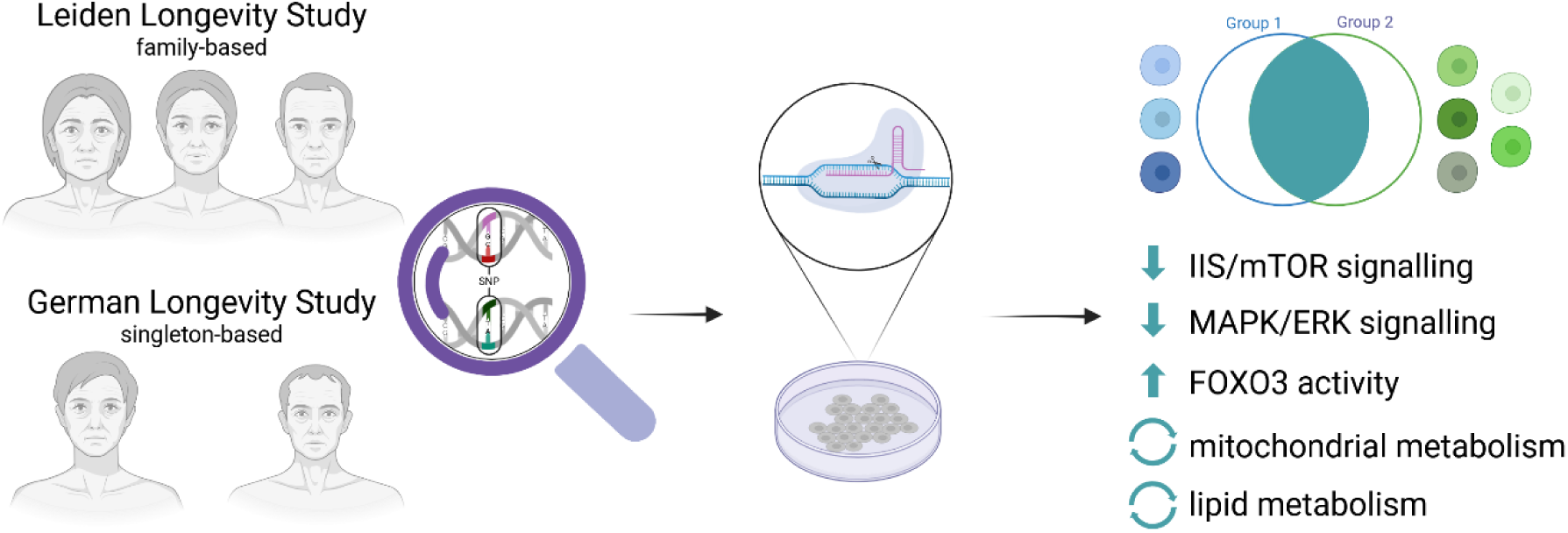

## Background

Human lifespan has increased substantially over the last two centuries (Oeppen and Vaupel 2002). However, healthspan, defined as the number of years in which an individual lives in relatively good health, without chronic illness (Kaeberlein 2018), did not show a proportional increase (Crimmins 2015). This discrepancy has resulted in a global rise in the number of individuals with multimorbidity (Chowdhury et al. 2023), thereby placing an increased burden on individuals, healthcare systems and the economy.

Exceptionally long-lived individuals and members of long-lived families exhibit compression of multimorbidity (Sebastiani et al. 2013; Andersen et al. 2012; Christensen et al. 2008; van den Berg et al. 2023) and can thus be used to identify mechanisms that can be targeted to extend both life- and healthspan. Longevity, the ability to survive to an exceptional old age, is influenced by numerous factors, including socioeconomic status, environment, behaviour, and genetic predispositions (Partridge et al. 2018). The extent to which genetics influence lifespan was estimated to be between 10 and 25% (Herskind et al. 1996; Kaplanis et al. 2018; Melzer et al. 2020; Ruby et al. 2018). However, a recent study demonstrated that this number is much higher (∼50%) when excluding extrinsic mortality (Shenhar et al. 2026).

In the case of long-lived individuals, genetic factors appear to play a prominent role in the regulation of their longevity, particularly within families where multiple individuals survive into the top 10% of their birth cohort (van den Berg et al. 2019). Utilising data from the Leiden Longevity Study (LLS), we previously demonstrated that ancestral familial longevity is associated with a 13-year delayed onset of the first cardiometabolic disease, which is coupled to a lower genetic predisposition to coronary artery disease (van den Berg et al. 2023; Sant’Anna Barbosa Ferreira et al. 2025). These findings show that LLS families are optimal for the study of the genetics underlying both longevity and cardiometabolic disease resilience. A recent hypothesis-free genetic linkage analysis of these families identified multiple rare protein-altering genetic variants, including a functional variant in *CGAS*, that may be involved in this resilience (Putter et al. 2025).

Non-family-based studies of human longevity have relied on genome-wide association studies (GWAS), with the aim of identifying common genetic variants (minor allele frequency (MAF) ≥1%) by comparing long-lived cases to shorter (average)-lived controls. These GWAS identified a limited set of robust genes that are associated with the regulation of longevity, i.e., *APOE* and *FOXO3* (Smulders and Deelen 2023). Consequently, the field has shifted towards the investigation of the impact of rare genetic variants (MAF <1%), a strategy that has already been applied for several other complex diseases and phenotypes (Marouli et al. 2017; Manolio et al. 2009). Studies in the field of ageing and longevity with a particular focus on rare genetic variants have found that the combined genetic variation within the insulin/insulin-like growth factor 1 signalling (IIS) and mechanistic target of rapamycin (mTOR) signalling pathway is associated with longevity (Lin et al. 2021; Kolbe et al. 2025). However, they did not investigate ancestral (more heritable) familial longevity and were limited to *in silico* analyses. We therefore previously utilised the LLS to investigate the genetics of ancestral familial longevity using a candidate longevity pathway approach, focussed on the mitogen-activated protein kinase/extracellular signal-regulated kinase (MAPK/ERK) signalling pathway. We identified protein altering genetic variants in *RAF1* and *NF1* that showed functional effects on the MAPK/ERK signalling activity, as well as general cellular health (Baghdadi et al. 2025). In the present study, we extended this approach to rare genetic variants in the IIS/mTOR signalling pathway. Moreover, we added variants in these pathways identified in an additional cohort consisting of individuals without a known familial history of longevity, the German Longevity Study (GLS) (Torres et al. 2021).

The IIS/mTOR signalling pathway represents a pivotal signalling hub that plays a crucial role in the regulation of growth, metabolism, and cellular survival (Burchfield et al. 2025). Moreover, the manipulation of this signalling network has proven its importance in the regulation of lifespan across species (Fontana et al. 2010). Genetic manipulation, for example through the (partial) knock-out of IRS1, IGF1R, and p70 S6K, or their orthologs, has been demonstrated to extend lifespan in worms, flies, and mice (Baghdadi et al. 2023; Clancy et al. 2001; Selman et al. 2008; Selman et al. 2011; Selman et al. 2009; Zhang et al. 2024; Kenyon et al. 1993; Holzenberger et al. 2003). Comparable results are obtained through nutritional (i.e., dietary restriction) and pharmacological (i.e. rapamycin, trametinib, lithium, and combinations thereof) inhibition of the IIS/mTOR and the connected MAPK/ERK signalling pathways (Gkioni et al. 2025; Slack et al. 2015; Di Francesco et al. 2024; Baghdadi et al. 2024; Miller et al. 2014; Castillo-Quan et al. 2019). The role of the IIS/mTOR signalling pathway in human longevity has thus far been studied by identifying single genetic variants in specific genes for which the orthologs have been implicated in lifespan regulation in model organisms, such as *FOXO3*, *IGF1*, and *IGF1R* (Tazearslan et al. 2011; Flachsbart et al. 2017; Willcox et al. 2008; Ali et al. 2025; Suh et al. 2008), or by examining the combined common or rare variation within the pathway (Deelen et al. 2011; Kolbe et al. 2025; Lin et al. 2021; Passtoors et al. 2013). These studies provide substantial evidence for the significance of the IIS/mTOR signalling pathway in human longevity. However, only limited functional validation for common variants was performed (Tazearslan et al. 2011) and this is lacking for rare variants.

In contrast to common genetic variants, there is insufficient statistical power to significantly associate single rare genetic variants with an outcome, unless one has access to a very large number of human samples, which is not the case for human longevity. Therefore, it is vital to perform functional validation experiments using *in vitro* or *in vivo* methods (Baghdadi et al. 2022). In this study, we perform *in vitro* characterisation of rare genetic variants in genes that play a role in the IIS/mTOR signalling pathway identified in the LLS. We further expanded the set of variants under investigation by incorporating genetic variants from the GLS for genes in which we had already identified multiple variants in the LLS.

We utilised CRISPR/Cas9 gene editing using the Cas9^D10A^ nickase enzyme (Cas9n) to generate transgenic mouse embryonic stem cell (mESC) lines. We additionally included mESC lines harbouring variants in the MAPK/ERK signalling pathway, which we identified in our previous work (Baghdadi et al. 2025). The basic functional characterisation of pathway activity leaned on this previous work and was extended by additional variants (in another pathway) and more in-depth characterisation. Our main aim was to identify shared mechanisms of the identified rare genetic variants that could be linked to the longevity observed in their carriers. In order to identify such mechanisms, we performed untargeted transcriptomics and proteomics. Furthermore, we evaluated the effects of the variants on the IIS/mTOR and MAPK/ERK signalling pathways, proliferation, and FOXO3 signalling. In light of the findings from the omics and informed by the literature on cellular ageing models, we performed lipidomics, stress assays, mitochondrial DNA (mtDNA) measurements and the Seahorse XF Cell Mito Stress Test. Given that we utilised a stem cell line, we also assessed the potential of the cells for spontaneous differentiation. While the cells clustered into two distinct groups for a subset of experimental readouts, they frequently exhibited the same phenotype for all variants studied, hinting at shared functional effects on cellular read-outs associated with organismal longevity in animal studies that may potentially be linked to the longevity of the human carriers.

## Materials and Methods

### Study population and selection criteria for genetic variants

In this study, we used genetic data from two studies: the Leiden Longevity Study (LLS) (Schoenmaker et al. 2006) and the German Longevity Study (GLS) (Kolbe et al. 2025). The LLS consists of Dutch sibships with at least two long-lived siblings. A description of the LLS and its inclusion criteria can be found elsewhere (Schoenmaker et al. 2006). The filtering criteria we applied here to identify promising rare genetic variants are the same as we described previously (Baghdadi et al. 2025). Except, here we filtered for variants in the IIS/mTOR signalling pathway. Additionally, in contrast to our previous work, the Sequenom MassARRAY system using iPLEX Gold genotyping assays (Sequenom, San Diego, CA, USA) was used for validation of variants.

To identify additional variants, next to those in the LLS, we also examined data from the GLS. The GLS is comprised of 1,265 unrelated German male and female exceptionally long-lived individuals between the ages of 94 and 110, as well as 4,195 geographically matched younger controls. Longevity was defined as passing the 95^th^ age-at-death percentile of their respective birth cohort (Kolbe et al. 2025). For the filtering of variant identified in the exome-sequencing of the GLS, we applied criteria similar to those for the LLS:

1. The variant resides within a gene in which we identified a variant in the LLS.
2. The variant is predicted to be protein-altering, defined by a Combined Annotation Dependent Depletion (CADD; version GRCh38-v1.6) score ≥ 20 (Schubach et al. 2024).
3. The variant is absent (MAF = 0%) in the GLS reference population and is either absent or has a very low frequency (MAF < 0.01%) in the general population, based on the gnomAD non-Finnish European reference population (Chen et al. 2024).
4. The variant was validated using Sanger sequencing.

### Culturing of mESCs

AN3-12 mESCs were generated previously (Elling et al. 2017), and were provided to us as a kind gift from Martin Denzel. Culturing conditions have been described previously (Baghdadi et al. 2025).

### Generation of variant mESC lines with CRISPR/Cas9n

Cloning, transfection, fluorescence-activated cell sorting (FACS), and genotyping for the generation of variant mESC lines with CRISPR/Cas9n was done as described previously (Baghdadi et al. 2025). In short, two guide RNAs and one repair template carrying the variant, as well as a silent novel restriction site and silent mutations preventing recutting, were designed per variant. The nickase mutant Cas9n was used to introduce two single-strand cuts in close proximity to the variant site. Guide RNAs (gRNAs) (**Table S1**), single-stranded DNA oligonucleotides (ssODNs) (**Table S2**), and primers (**Table S3**) used here, can be found in the supplementary information. The generated mESC lines used for experiments were diploid and carried the variants in a homozygous state.

### Plating cells for transcriptomics and proteomics

Transcriptomics and proteomics were performed on each four replicates for 11 mESC lines: the wild-type and lines harbouring the following mutations: DEPTOR^Cys102Trp^, IRS1^Pro4Thr^, IRS1^Tyr465His^, PHLPP1^Pro298Leu^, PHLPP1^Leu843Pro^, PHLPP1^Glu1149Gly^, NF1^Phe1110Leu^, and RAF1^Asp633Tyr^. For the IRS1^Pro4Thr^ and NF1^Phe1110Leu^ variants, we used two independent cell lines each. To generate the replicates, cells were plated onto two 10 cm culture dishes per cell lines and were treated as independent lines. At passage 2 and 4, each two 100 mm cell culture dishes per omics measurement were plated with 1.25 million cells per dish. After 18 hrs RNA and protein samples were isolated.

### Transcriptomics

#### RNA extraction and data generation

Total RNA was isolated using 900 µL of TRIzol reagent (Invitrogen, CA, USA) according to the manufacturer’s instructions. GlycoBlue Coprecipitant (Thermo Fisher Scientific, MA, USA) was used to stain the RNA precipitate. RNA pellets were dissolved in 30 µL RNase-free water. The concentration and quality of the RNA were measured using TapeStation (Agilent Technologies, California, USA) and NanoDrop (Thermo Fisher Scientific). Ribosomal RNA removal, library preparation, and RNA-sequencing were performed by the Cologne Centre for Genomics. Libraries were sequenced as 100 bp paired-end reads with a depth of approximately 40 million reads per sample.

#### Transcriptomic data analysis

The transcriptomics data were processed and analysed with a custom-made pipeline using R packages. In short, raw reads were trimmed using *Trimmomatic*; *FastQC* and *MultiQC* were used for read quality control. Transcript counts were estimated and summarized at gene level using R packages *Kallisto* and *tximport*, respectively. Principal component analysis (PCA) was performed on variance-stabilized transformed (VST) expression data, computed using *DESeq2*’s VST function. The 500 most variable genes were selected by ranking on across-sample variance, and PCA was computed using R’s prcomp function. Samples were visualized along the first two principal components, and 95 % confidence ellipses were drawn to highlight predefined cell states. Differential expression analysis was performed using *DESeq2*. Low-count genes were excluded prior to fitting (retaining genes with ≥10 counts in at least 2 samples). Log2 fold changes were shrunk using the apeglm method. The R package *clusterProfiler* was used for overrepresentation analysis (ORA) with pathways from the MSigDB database. Plots were generated using base R, and the R package *ggplot2*. Variant cell lines were tested against the wild-type either individually or grouped together based on the PCA results. In case of the comparison per group, samples were first grouped together and the resulting groups were then jointly analysed.

### Proteomics

#### Protein digestion and peptide cleaning

To isolate proteins, cells were pelleted, washed twice with cold PBS, and stored as dry pellets at -80 °C. Cells were dissolved in 30 μL lysis buffer (6 M GuHCI, 2.5 mM TCEP, 10 mM CAA, and 100 mM Tris-HCl). For lysis, samples were heated at 95 °C for 10 min and afterwards sonicated using a Bioruptor (Diagenode, Belgium) at the following setting: 30 sec sonication, 30 sec break, 10 cycles, high performance at 4 °C. Samples were centrifuged at 20,000 g for 20 min. Supernatant containing 100 μg of protein was diluted 10x with 20 mM Tris, and digested with trypsin 1:200 (w/w) overnight. The digestion was stopped by adding 50% formic acid to a final concentration of 1% formic acid, and samples were cleared by centrifugation at 20,000 g for 10 min.

Peptide cleaning was performed with in-house 30 μg C18-SD StageTips. In brief, these tips were wetted with methanol, then 40% acetonitrile/0.1% formic acid and equilibrated with 0.1% formic acid by centrifugations of 1-2 min, without letting the tips dry in between these steps. The supernatant containing the total protein digest was loaded in 0.1% formic acid and 2x washed with 0.1% formic acid. To elute the peptides, 100 μL 40% acetonitrile/0.1% formic acid were added at 300 g for 4 min, and the peptides were dried in a SpeedVac at 45 °C for 45 min. The peptides were resuspended in 20 μL 0.1% FA, and concentrations were measured with a NanoDrop spectrophotometer (Thermo Fisher Scientific).

Further processing, measurements, and data pre-processing were performed by the Proteomics Core Facility of the Max Planck Institute for Biology of Ageing (MPI-Age). Protein extracts were adjusted to a concentration of ∼10 mg/mL in 6 M guanidinium chloride in 100 mM Tris-HCl. 10 μL of each of these initial protein extracts was introduced into a Kingfisher DW 96-well plate (Thermo Fisher Scientific). Immediately before further processing, these extracts were diluted 10:1 through the addition of 90 µL of Tris buffer and the plate was placed in the Kingfisher APEX (Thermo Fisher Scientific).

200 µg of MagReSyn Hydroxyl beads (ResynBio, South Africa) were robotically added to each well and mixed with the diluted protein solution. The plate was then removed from the robot and PAC was initiated by the addition of pure ethanol to a concentration of 70%. The plate was returned to the robot for mixing during the precipitation process. The beads with bound precipitated protein were then washed 3 times in 80% ethanol. They were then transferred to an elution plate, where they were incubated for 6 hrs in 50 mM ABC buffer (pH 7.8) with 1 µg of Trypsin Gold (Promega, WI, USA) under gentle mixing at 37 °C. Following this elution, beads were left overnight in the trypsin solution at 12 °C. The next day, the beads were eluted again in 0.1% TFA, and the two elutions were pooled.

#### Liquid chromatography

The Evosep One liquid chromatography system was used for analysing the samples with the 30 samples per day (30SPD) method. The analytical column we used was a ReproSil-Pur column, 15 cm x 150 µm, with 1.5 µm C18 beads (EV1137 Performance Column, Evosep, Denmark). The mobile phases A and B were 0.1 % formic acid in water and 0.1% formic acid in 100% ACN, respectively.

#### Mass spectrometry

Peptides were analysed on a hybrid TIMS quadrupole TOF mass spectrometer (timsTOF HT, Bruker, MA, USA) in a data-independent acquisition parallel accumulation, serial fragmentation (diaPASEF) mode. The mass spectra range was set to 350 – 1,200 m/z and TIMS ion accumulation and ramp times were set to either 50 ms or 100 ms, and total cycle time was 0.73s or 1.38s. The ion mobility range was set to 1/K0 = 0.8 - 1.25 V-s/cm^2^. Isolation windows in the m/z versus ion mobility plane were defined to cover the region of highest precursor ion density, using py_DIA. Collision energy was applied linearly with ion mobility from 0.6 to 1.6 V-s/cm^2^, and collision energy from 20 to 59 eV.

#### Proteomic data analysis

Raw data were analysed using Spectronaut version 20.0.25 (Biognosys, Switzerland), using the default parameters against the one-protein-per-gene reference proteome for *Mus musculus*, UP000000589, downloaded August, 2022. Methionine oxidation and protein N-terminal acetylation were set as variable modifications; cysteine carbamidomethylation was set as fixed modification. The digestion parameters were set to “specific” and “Trypsin/P,” with two missed cleavages permitted. Protein groups were filtered for at least two valid values in at least one comparison group, and missing values were imputed from a normal distribution with a down-shift of 1.8 and standard deviation of 0.3.

The proteomics data were processed and analysed with a custom-made pipeline using R packages. PCA was performed on the 500 most variable proteins, selected by ranking features on their across-sample variance. Abundance values were mean-centered and scaled prior to PCA. PCA was computed using the prcomp function in R, and samples were visualized along the first two principal components. 95% confidence ellipses were drawn to highlight predefined cell state groups. Differential abundance analysis was performed using the R package *limma*. The R package *clusterProfiler* was used for ORA with pathways from the MSigDB database. Plots were generated using base R, and the R-package *ggplot2*. Variant cell lines were tested against the wild-type either individually or grouped together based on the PCA results

### Lipidomics

#### Lipid isolation

Lipidomics were performed on the same 11 mESC lines as for transcriptomics and proteomics. Samples were taken at two different passages. Per passage, 1 million cells per cell line were collected in triplicate. Cell pellets were washed twice with PBS. Washed pellets were frozen at -70 °C.

Extraction buffer was prepared freshly in a 3:1 [v:v] mixture of Methyl tert-butyl ether (Sigma-Aldrich, Mo, USA), Optima LC-MS grade methanol (Thermo Fisher Scientific). Per 100 mL extraction buffer the following standards are added: 10 µL 2.5 mM U-^13^C^15^N amino acids, 10 µL of 1 mg/mL ^13^C_10_ ATP, 20 µL of 100 µg/mL citric acid D4, 20 µL EquiSPLASH™ LIPIDOMIX (Avanti Polar lipids), and 20 µL of 1 mg/mL cholesterol D7. For metabolite extraction sample tubes were placed on dry ice and 1 mL of pre-cooled extraction buffer was added per sample. Samples were sonicated at 85% intensity three to five times with a tip sonicator (Sonics, CT, USA), followed by a 30 min incubation at 4 °C and 1,500 rpm in the thermomixer. Supernatant was collected after centrifugation at 21,000 g and 4 °C for 10 min. 500 µL of H_2_0:methanol 3:1 [v:v] was added and incubated on an orbital mixer at 1,500 rpm and 15 °C for 10 min. Samples were phase separated by centrifugation at 5,000 g and 15 °C for 3 min. 700 µL of the upper lipid phase were transferred to a new tube. Samples were dried down in a SpeedVac concentrator at 1,000 rpm and 20 °C and frozen at -80 °C until measurement.

#### Liquid Chromatography-High Resolution Mass Spectrometry-based analysis of lipids

Measurement of lipids as well as preprocessing of the data were performed by the Metabolomics Core Facility of the MPI-Age. The stored (−80 °C) lipid extracts were re-suspended in 150 µL of a UPLC-grade acetonitrile: isopropanol (70:30 [v:v]) mixture, followed by vortexing and 10 min incubation on a thermomixer at 4°C. The re-suspended samples were cleared by a 5 min centrifugation at 16,000 g and the supernatants were transferred to 2 mL glass vials with 300 µL glass inserts (Chromatography Zubehör Trott, Germany). From each analytical sample a volume of 20 µL was mixed together to generate a sample pool. These pools were used as instrumental and sample stability quality controls, which are run after every 10^th^ analytical sample in the sample sequence, or after each replicate group.

All samples and pools were placed in a Vanquish UHPLC chromatography system, equipped with a quaternary pump (Thermo Fisher Scientific), and kept in the sample manager at 6 °C. The UHPLC was connected to a timsTOF Pro 2 HRMS, equipped with a heated ESI (VIP-HESI) source (Bruker).

Of each lipid sample, 1 µL was injected on a premier CSH C18 100 x 2.1 mm UPLC column, packed with 1.7 µm particles (Waters, MA, USA) kept at 50 °C. The flow rate of the UPLC was set to 400 µL/min, and the buffer system consisted of buffer A (10 mM ammonium formate, 0.1% formic acid in UPLC-grade acetonitrile:water (60:40, [v:v])) and buffer B (10 mM ammonium formate, 0.1% formic acid in UPLC-grade isopropanol:acetonitrile (90:10, [v:v])). The UPLC gradient was as follows: 0-0.5 min 45-48% B, 0.5-1 min 48-55% B, 1-1.8 min 55-60% B, 1.8-10 min 60-85% B, 10-11 min 85-99% B, 11-11.5 min 99% B, and 11.7-15 min re-equilibrating at 45% B. This leads to a total runtime of 15 min per sample.

At the beginning of each run, the mass accuracy and mobility were recalibrated by injection of a mixture of 10 mM sodium formate calibrant solution and ESI-L Low Concentration Tuning Mix (Agilent) (1:1 v/v). Data-dependent parallel accumulation serial fragmentation acquisition mode was used. Main analysis was run in positive ionization mode. Source parameters were as follows. Capillary voltage was set to 4’500 V with an end plate offset of 500 V. Nebulizer pressure was set to 2 bar, dry gas was running with 8 L/min and a temperature of 230 °C. Sheet gas was running with 4 L/min and a temperature of 400 °C. For MS2 experiments, masses were isolated with a width of 2 Da and fragmentation was induced with a collision energy of 30 eV. To gain more information about fatty acid composition of lipids several injections of pooled quality controls were performed in negative mode. For negative mode collision energy was set to 40 eV and capillary voltage was set to -3,500 V.

#### Lipidomic analysis

Raw data were further processed with MetaboScape (v. 2024) to generate a feature table and to annotate lipid species. Lipid species annotation was further validated manually. Moreover, only lipid species that were 3 times higher in the quality controls samples as compared to the method blank samples were kept for further analysis. For PCA, missing values were imputed, and batch corrected for passage using the R package *limma*. PCA was performed on the batch-corrected data matrix using prcomp in R. Samples were visualized along the first two principal components, with 95% confidence ellipses drawn to highlight predefined cell state groups. Linear modelling for individual lipid species was done on log transformed and median normalised data. The median was taken per lipid class for representation of data in a heatmap. Results are given per variant group as Log2 fold change compared to the wild-type mESC line. R packages, *ggplot2*, *grid*, and *ggthemes* were used to visualise the results of the analyses.

#### Protein isolation and western blotting

Protein isolation of normal state and insulin stimulated samples and western blotting were performed as described previously (Baghdadi et al. 2025). The only deviations from this protocol were that 400,000 cells/well were plated in a 6-well plate, and GAPDH was used as the housekeeping protein. The primary antibodies used are depicted in **Table S4**.

#### RNA isolation and qPCRs

For RNA extraction 400,000 cells were plated per well in a 6-well plate. On the next day total RNA isolation was started using 300 µL TRIzol Reagent (Invitrogen) per well, according to manufacturer’s instructions. DNA-free DNA Removal Kit (Invitrogen) was used according to manufacturer’s instructions. The quantity and quality of RNA were assessed using a NanoDrop spectrophotometer (Thermo Fisher Scientific). The SuperScript III First-Strand Synthesis SuperMix (Invitrogen) was used according to manufacturer’s instructions for cDNA generation using 5,000 ng of RNA. Remaining RNA was stored at -70 °C. cDNA was diluted 1:4 for qRT-PCR. TaqMan Assay probes (**Table S5)** and the TaqMan Gene Expression Master Mix (Thermo Fisher Scientific) were used. Samples were loaded in quadruplicates in 384 well plates on the QuantStudio 6 Flex Real-Time PCR System (4485691, Thermo Fisher Scientific) with a reaction size of 10 µL, a standard curve, and the standard speed. Gene expression was calculated by ΔΔCt after normalisation to the reference gene *Gapdh*.

#### FOXO3 binding assay

3,000 cells/well were plated in 100 µL complete growth medium in a white 96-well plate. 6 hrs after plating, cells were transfected with 0.125 µg of DNA per well, using Lipofectamine 3000. The two plasmids Forkhead responsive element (FHRE)-luc (Addgene: #1789) and pGL4.75 (kind gift from Adam Antebi; Promega) were used in a ratio of 1:1.21 to account for equal copy numbers. The Renilla luciferase plasmid pGL4.75 was used to control for transfection efficiency and cell number. 48 hrs after transfection, medium was exchanged with 30 uL PBS per well. The Dual-Glo® Luciferase Assay System (Promega) was used according to manufacturer’s instructions with a volume of 30 uL per reagent. The firefly luminescence (FHRE-luc) was normalised to the Renilla luminescence (pGL4.75) per well. The normalised measurements are shown as a ratio to the average of the wild-type.

#### Growth assessment

Growth rate was assessed using the Incucyte Live-Cell Analysis system (Sartorius, Germany) as described previously (Baghdadi et al. 2025). In short, increase in confluence over time was measured using the Incucyte.

#### Differentiation

The spontaneous differentiation potential was assessed by qPCR. Cells were plated in complete growth medium in 6-well plates (**Table 1**). 6 hrs after plating, cells were washed twice with PBS and medium was changed to growth medium without leukaemia inhibitory factor (LIF) (-LIF medium). For the control (Day 0) the medium was not changed. RNA was isolated after 1, 3, and 7 days of growth. For the day 7 samples -LIF medium was refreshed at day 3 and day 6 after plating.

**Table 1.**
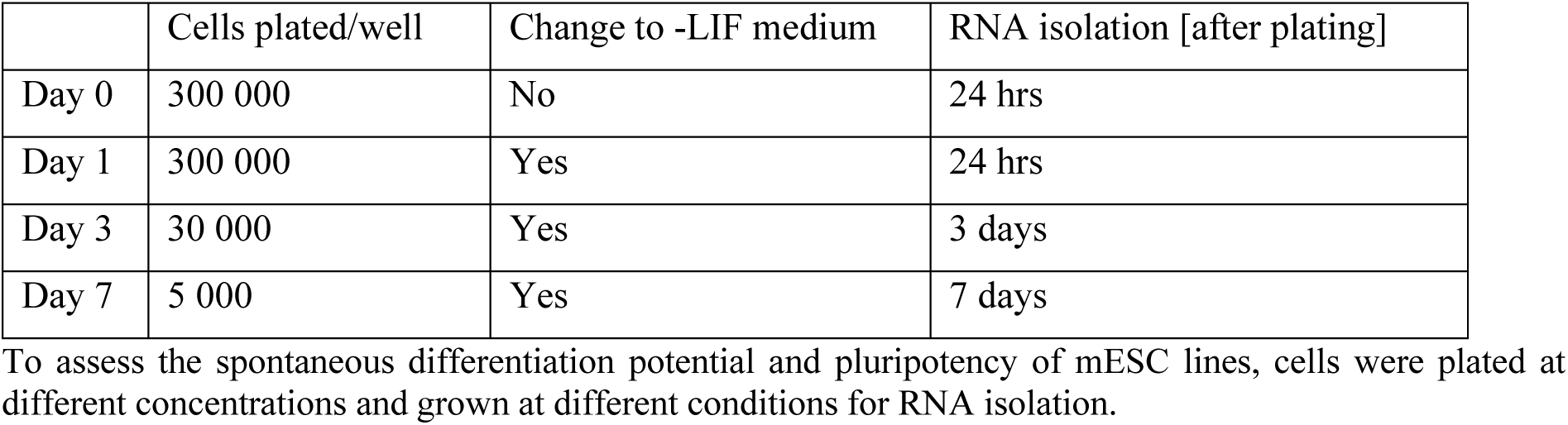
Plating for differentiation experiment.

|  | Cells plated/well | Change to -LIF medium | RNA isolation [after plating] |
| --- | --- | --- | --- |
| Day 0 | 300 000 | No | 24 hrs |
| Day 1 | 300 000 | Yes | 24 hrs |
| Day 3 | 30 000 | Yes | 3 days |
| Day 7 | 5 000 | Yes | 7 days |

To assess the spontaneous differentiation potential and pluripotency of mESC lines, cells were plated at different concentrations and grown at different conditions for RNA isolation.

For RNA isolation cells were detached from the plates in 300 µL Trizol and frozen at -70 °C. RNA isolation, cDNA generation, and qPCRs were performed as described above, with the deviation that *Power* SYBR Green PCR Master Mix (Applied Biosystems, MA, USA) was used. Primer sequences were taken from Kroef and colleagues (Kroef et al. 2025) and *Rpl37a* was used as a housekeeping gene.

#### Stress assays

The resistance to replicative stress was assessed as previously described (Baghdadi et al. 2025). In short, the confluence of mESCs exposed to stressors at different concentrations was measured 22 hrs after the addition of stressors, using the Incucyte. Per mESC line, confluence was normalised to the respective cell line without the addition of stressors. Additionally, propidium iodide was used at a final concentration of 1 µg/mL to stain dead cells. The three stressors used here were all diluted in water. Cobalt chloride (C8661, Sigma-Aldrich) and hydroxyurea (H8627, Sigma-Aldrich) were freshly diluted before every experiment, while for cadmium chloride (C3141, Sigma-Aldrich) a stock was prepared and frozen. The final concentrations used were: cobalt chloride: 0, 100, 200, 800 µM; hydroxyurea: 0, 0.43, 1.73 and 4.32 mM; cadmium chloride: 0, 2, 6 and 12 µM. Propidium iodide positive cells were identified using the segmentation “Surface Fit” and a “Threshold (RCU)” of 1. For cell death analysis, per well, the propidium iodide count was divided by the confluence. This cell death ratio was normalised to the control without stressor of the respective mESC line.

#### Seahorse

To assess mitochondrial function the Seahorse XF Cell Mito Stress Test and a Seahorse XFe96 Analyzer (Agilent Technologies) were used according to manufacturer’s instructions. Cells were plated at previously determined concentrations (24,000 cells/well for wild-type, Deptor^Cys102Trp^, Raf1^Asp633Tyr^, and Phlpp1^Leu843Pro^; 18,000 cells/well for Phlpp1^Pro298Leu^ and Phlpp1^Glu1149Gly^) to assure a comparable confluence at the time of measurement. 24 hrs after plating, cell medium was changed to Seahorse assay medium (XF Base with 1 mM sodium pyruvate, 2 mM L-glutamine, 10 mM glucose (Agilent Technologies) and 0.01% LIF (Millipore)). The following inhibitors were used: 1 µM Oligomycin, 0.5 µM FCCP and 0.5 µM Rotenone/Antimycin A. All data were normalised to protein concentration measured by a BCA assay (Thermo Fisher Scientific) immediately after completion of the assay.

#### Measurement of mtDNA levels by qPCR

To measure mtDNA levels, the NucleoSpin Tissue XS kit (Macherey-Nagel, Germany) was used to isolate whole DNA from mESCs. 14 ng of total DNA was used per well in a Taq Man qPCR (details described above). TaqMan probes against *Cox1* and *Rnr2* were used as mitochondrial probes. *18S* was used as a genomic DNA probe. TaqMan probes used can be found in **Table S5**. Per sample, mean Cts of each mitochondrial probe were normalised to the *18S* mean Ct of the respective sample. Mt/nuclear DNA probe ratios were normalised to the mean of the wild-type.

#### Statistics

Unless stated differently, data were analysed using GraphPad Prism 11.0.0. Significance levels are indicated as \**P* < 0.05, \*\**P* < 0.01, and \*\*\**P* < 0.001. Across group comparisons were made using a one-way or two-way ANOVA and between groups using a Dunnett’s post hoc test. Error bars correspond to standard deviations.

#### AI Tool Usage

During the preparation of this article, the authors used an AI language model to assist with editing and revision of the text in regards to grammar, spelling, and phrasing. The AI tool was used solely for language refinement. All scientific content, data interpretation, and final manuscript approval were performed by the authors. The AI tool did not contribute to the conception, design, analysis, or interpretation of the study.

## Results

### Identification of rare protein-altering genetic variants in exceptionally long-lived individuals

In order to investigate the functional effects of rare protein-altering genetic variants that may contribute to human longevity, we employed a targeted pathway approach focussing on the IIS/mTOR signalling pathway, which has shown to play a vital role in the regulation of lifespan in model organism-based studies. We used the family-based LLS and the singleton-based GLS for the selection of variants. The filtering criteria we applied to identify the most promising variants were introduced before (Baghdadi et al. 2025), with some minor adjustments (see Methods section). In the current study, we focussed on the IIS/mTOR signalling pathway. In addition, we selected variants from two populations (LLS + GLS) instead of one (LLS). We identified an initial set of rare variants that were carried by at least two long-lived siblings in one family in the LLS (**Figure 1**) and expanded the list with variants from the GLS residing within the same genes in which we identified an LLS variant of interest. For *PHLPP1* and *IRS1*, we identified two and three additional rare genetic variants, respectively, in the GLS. All identified variants were carried in a heterozygous state by the exceptionally long-lived individuals. An overview of all variants identified in IIS/mTOR signalling in the LLS and the additional variants identified in the same genes in the GLS is depicted in **Table 2**.

**Figure 1.**
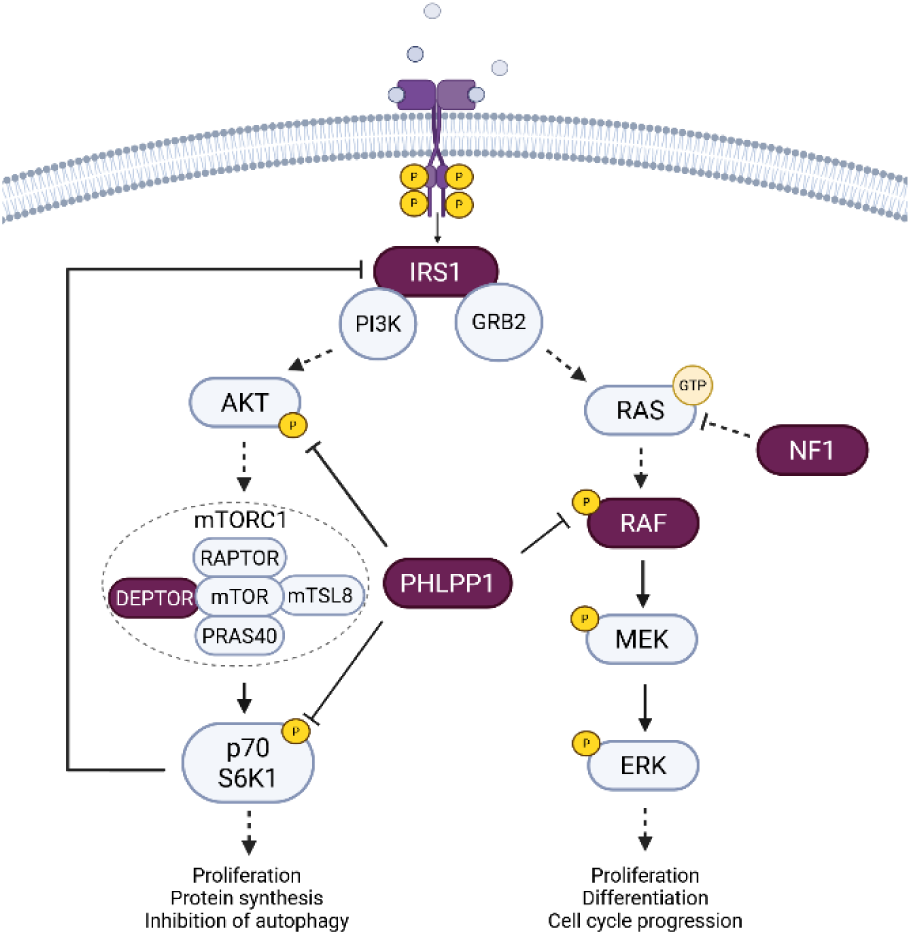
Rare genetic variants in the IIS/mTOR and MAPK/ERK signalling pathways. Simplified schematic of the IIS/mTOR (left arm) and MAPK/ERK (right arm) signalling pathways. Proteins in which variants were identified in this study are highlighted in purple.

**Table 2.**
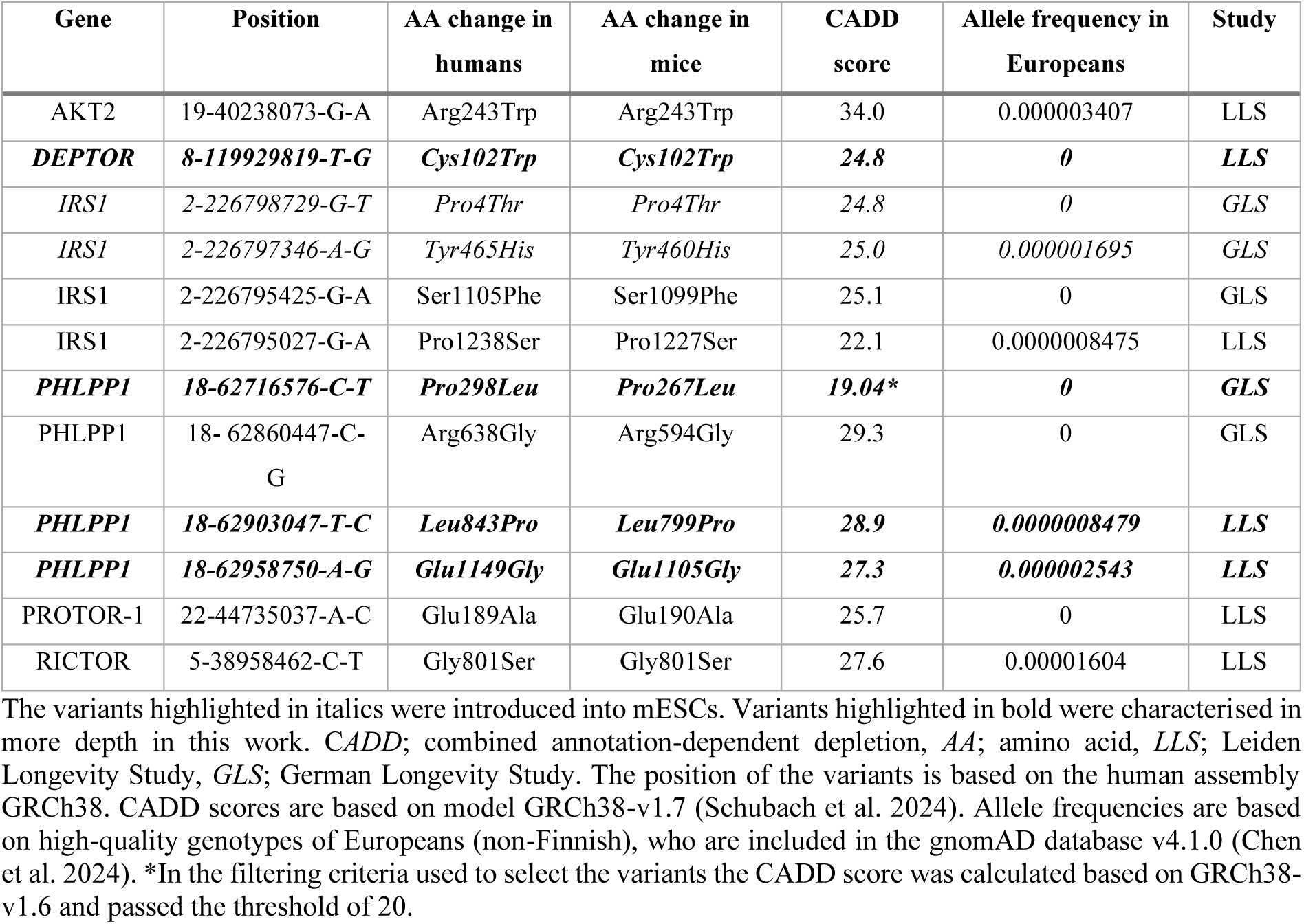
Rare genetic variants in the IIS/mTOR signalling pathway identified in exceptionally long-lived individuals from the Leiden Longevity Study and German Longevity Study.

| Gene | Position | AA change in humans | AA change in mice | CADD score | Allele frequency in Europeans | Study |
| --- | --- | --- | --- | --- | --- | --- |
| AKT2 | 19-40238073-G-A | Arg243Trp | Arg243Trp | 34.0 | 0.000003407 | LLS |
| **DEPTOR** | **8-119929819-T-G** | **Cys102Trp** | **Cys102Trp** | **24.8** | **0** | **LLS** |
| *IRS1* | 2-226798729-G-T | *Pro4Thr* | *Pro4Thr* | 24.8 | 0 | *GLS* |
| *IRS1* | 2-226797346-A-G | *Tyr465His* | *Tyr460His* | 25.0 | 0.000001695 | *GLS* |
| IRS1 | 2-226795425-G-A | Ser1105Phe | Ser1099Phe | 25.1 | 0 | GLS |
| IRS1 | 2-226795027-G-A | Pro1238Ser | Pro1227Ser | 22.1 | 0.0000008475 | LLS |
| **PHLPP1** | **18-62716576-C-T** | **Pro298Leu** | **Pro267Leu** | **19.04\*** | **0** | **GLS** |
| PHLPP1 | 18- 62860447-C-G | Arg638Gly | Arg594Gly | 29.3 | 0 | GLS |
| **PHLPP1** | **18-62903047-T-C** | **Leu843Pro** | **Leu799Pro** | **28.9** | **0.0000008479** | **LLS** |
| **PHLPP1** | **18-62958750-A-G** | **Glu1149Gly** | **Glu1105Gly** | **27.3** | **0.000002543** | **LLS** |
| PROTOR-1 | 22-44735037-A-C | Glu189Ala | Glu190Ala | 25.7 | 0 | LLS |
| RICTOR | 5-38958462-C-T | Gly801Ser | Gly801Ser | 27.6 | 0.00001604 | LLS |

The variants highlighted in italics were introduced into mESCs. Variants highlighted in bold were characterised in more depth in this work. CADD; combined annotation-dependent depletion, AA; amino acid, LLS; Leiden Longevity Study, GLS; German Longevity Study. The position of the variants is based on the human assembly GRCh38. CADD scores are based on model GRCh38-v1.7 (Schubach et al. 2024). Allele frequencies are based on high-quality genotypes of Europeans (non-Finnish), who are included in the gnomAD database v4.1.0 (Chen et al. 2024). \*In the filtering criteria used to select the variants the CADD score was calculated based on GRCh38-v1.6 and passed the threshold of 20.

In order to assess commonalties of different mechanisms of longevity and given the cross-talk between the IIS/mTOR and MAPK/ERK signalling pathways, we decided to additionally include the two variants, RAF1^Asp633Tyr^ and NF1^Phe1110Leu^ identified in our previous candidate pathway-based study (Baghdadi et al. 2025) (**Figure 1**).

For the purpose of *in vitro* functional characterisation we introduced the variants into the mESC line AN3-12 using CRISPR/Cas9 gene editing with the Cas9^D10A^ nickase enzyme (double-nicking strategy), applying the same protocol as in our previous study (Baghdadi et al. 2025). All variants were introduced into the AN3-12 cells in a homozygous state.

### Proteomic and transcriptomic analysis

In order to assess the variants’ broad effects on a molecular level, we performed normal growth state transcriptomic and proteomic measurements. PCA revealed a clustering of the variant mESC lines into two distinct groups at both the transcriptomic and proteomic level, with both of these groups demonstrating a clear separation from the wild-type mESCs (**Figure 2**) (Group 1: DEPTOR^Cys102Trp^, RAF1^Asp633Tyr^, and PHLPP1^Leu843Pro^; Group 2: IRS1^Pro4Thr^, IRS1^Tyr465His^, NF1^Phe1110Leu^, PHLPP1^Pro298Leu^, and PHLPP1^Glu1149Gly^). PC1 separates the two groups from each other, while PC2 separates all variants from the wild-type. We performed non-directional overrepresentation analysis (ORA) per PCA group against the wild-type using the MSigDB hallmark pathways. For the proteomics, the hallmark “Mtorc1 Signaling” was enriched in both groups. Group 1 demonstrated a specific enrichment for “Xenobiotic Metabolism” and Group 2 for “Glycolysis” and “Hypoxia”. In the case of the transcriptomics, a greater number of hallmarks were enriched. Among them were “Glycolysis”, “Hypoxia”, “Cholesterol Homeostasis”, and “Epithelial Mesenchymal Transition” for both groups. Furthermore, both groups demonstrated an effect on the hallmark “Kras Signaling Up”, while Group 2 additionally showed enrichment for the hallmark “Kras Signaling Dn”. In the transcriptomics dataset, *Phlpp1* and *Irs1* transcript levels were significantly downregulated in all variant mESC lines while *Nf1* transcript levels were increased for Group 1 variant mESCs and decreased for Group 2 variant mESCs (**Figure S1**). *Raf1* and *Deptor* transcript levels remained unaltered. This indicates that the variants may not necessarily affect the expression of the gene or protein in which they are residing but (additionally) possibly regulate downstream signalling through changes in posttranscriptional and posttranslational modifications.

**Figure 2.**
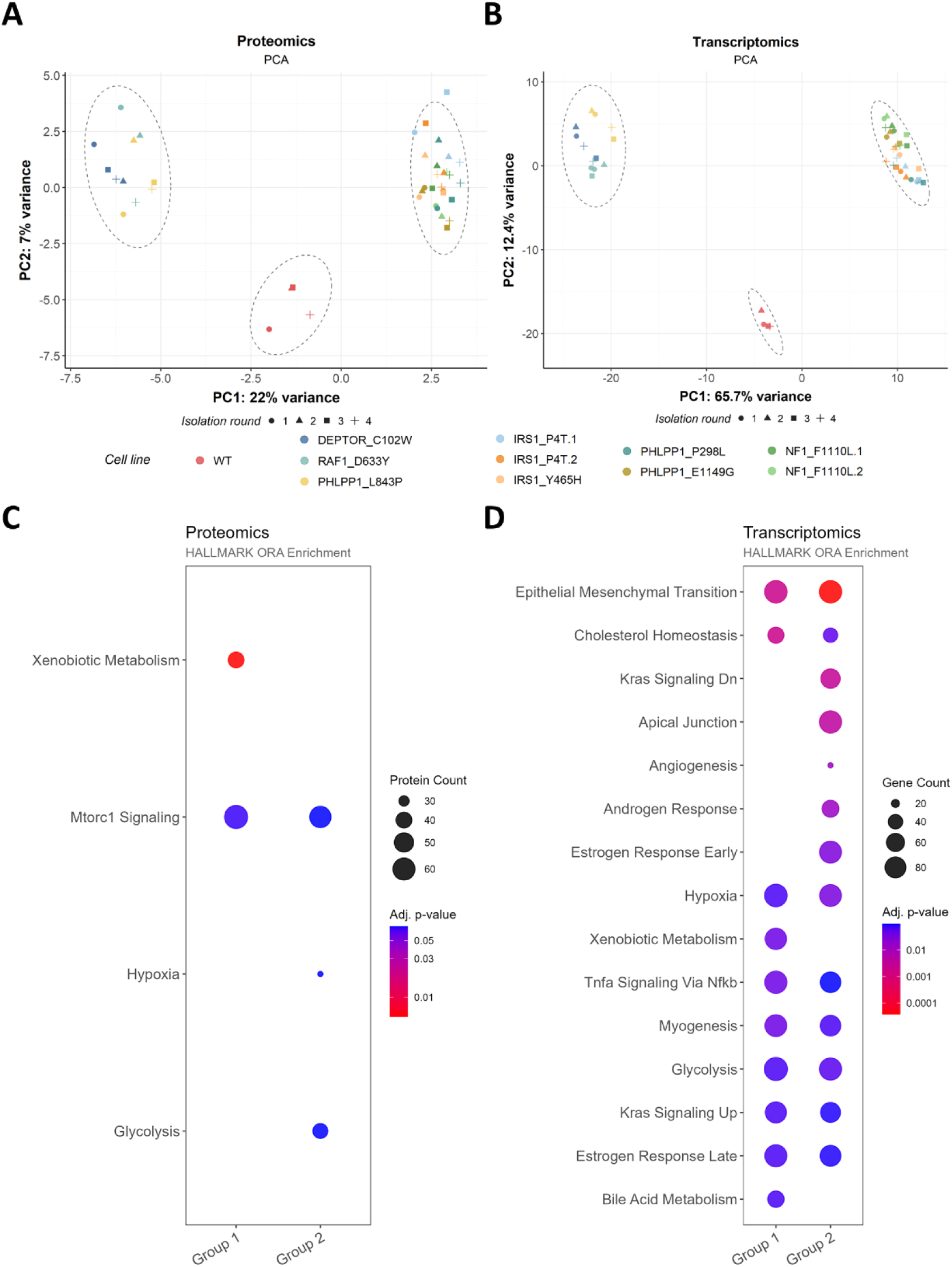
Matched proteomics and transcriptomics of variant mESC lines. PCA of proteomics (**A**) and transcriptomics (**B**) of the 500 most variable proteins/transcripts across all samples. For each cell line 4 replicates were isolated, represented by different symbols. ORA on significantly deregulated proteins (**C**)/transcripts (**D**) (p≤0.05) using MSigDB hallmarks. In the ORA, samples were grouped according to the clustering they show in the PCA (Group 1: DEPTOR^Cys102Trp^, RAF1^Asp633Tyr^, and PHLPP1^Leu843Pro^; Group 2: IRS1^Pro4Thr^, IRS1^Tyr465His^, NF1^Phe1110Leu^, PHLPP1^Pro298Leu^, and PHLPP1^Glu1149Gly^). *WT*; wild-type AN3-12 line.

Due to technical feasibility, we conducted the subsequent functional characterisation with five variant mESC lines and the wild-type. We selected all three variants that are part of Group 1 (i.e., DEPTOR^Cys102Trp^, RAF1^Asp633Tyr^, and PHLPP1^Leu843Pro^) as well as the two PHLPP1 variants that belong to Group 2 (PHLPP1^Pro298Leu^ and PHLPP1^Glu1149Gly^).

### The variant mESC lines show consistently lowered mTORC1 and MAPK/ERK signalling pathway activity

In order to evaluate the downstream regulation of the variants on IIS/mTOR and MAPK/ERK signalling pathway activity, we performed western blotting of normal state as well as insulin stimulated mESCs. The activity of both the IIS/mTOR and MAPK/ERK signalling pathways is regulated through post-translational modifications (Vargas-Ibarra et al. 2021; Yin et al. 2021). Therefore, the assessment of the level of a total protein and its respective phosphorylated version, for example through western blotting, can be used to determine the pathway activity.

We examined the phosphorylation of p70 S6K (Thr389) and AKT (Thr308), both readouts of mTORC1 activity, as well as the phosphorylation of AKT (Ser473), which is a readout of mTORC2 activity (Sarbassov et al. 2005). Phosphorylation of AKT (Ser473) was upregulated in all variant mESC lines in the normal state (**Figure 3A**). This finding is consistent with our previous observations for the RAF1^Asp633Tyr^ and NF1^Phe1110Leu^ cell lines (Baghdadi et al. 2025) and suggests increased mTORC2 activity under normal growth conditions. Furthermore, the upregulation of phosphorylation at AKT (Ser473) is in line with the downregulation of *Phlpp1* transcripts in both variant groups, as PHLPP1 is known to dephosphorylate AKT at Ser473 (Gao et al. 2005).

**Figure 3.**
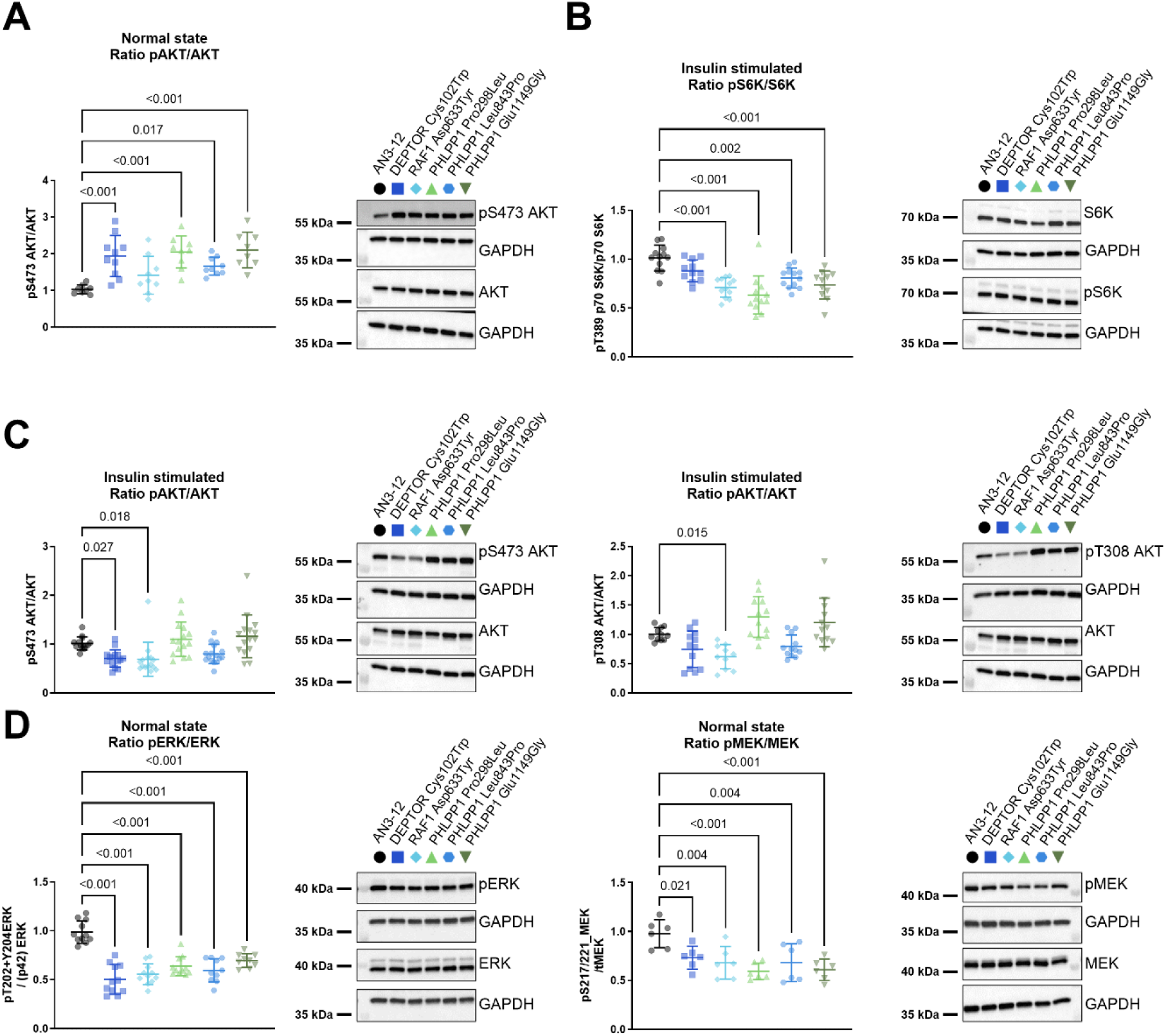
Variants alter activity of IIS/mTOR and MAPK/ERK signalling pathways. Analysis of IIS/mTOR (**A**-**C**) and MAPK/ERK (**D**) signalling pathways by measuring phosphorylation ratios through western blotting. Normal state (**A**, **D**) phosphorylation was assessed in mESCs grown in complete growth medium. For insulin stimulation (**B**, **C**) mESCs were serum-starved for 6 hrs before a 10 min insulin stimulation (100 nM). **A** Increase of phosphorylated AKT (Ser473) over total AKT in all variant mESC lines under normal growth conditions. **B** Decrease of phosphorylated p70 S6K (Thr389) over total p70 S6K in all variant mESC lines after insulin stimulation. **C** Decrease of phosphorylated AKT (Ser473) and AKT (Thr308) over total AKT, respectively, for mESC lines harbouring DEPTOR^Cys102Trp^, RAF1^Asp633Tyr^ and PHLPP1^Leu843Pro^ (Group 1) after insulin stimulation. **D** Decrease in phosphorylation of ERK1/2 (Thr202+Tyr204) over total ERK1/2 and MEK1/2 (Ser217/221) over total MEK1/2 in all variant mESC lines under normal growth conditions. For all experiments, GAPDH was used for normalisation. Data shown are from two to three independent experiments with four technical replicates each. Error bars represent standard deviations. Data were analysed using a one-way ANOVA and Dunnett’s post hoc test. The quantified data of the phosphorylated and total proteins are provided in **Figure S3**.

To stimulate the IIS/mTOR signalling pathway, we serum starved the cells for six hrs followed by stimulation with 100 nM insulin for 10 min. Following stimulation, the level of phosphorylation of p70 S6K (Thr389) was decreased in all variant mESC lines compared to the wild-type (**Figure 3B**), while the level of phosphorylation of AKT (Ser473) was reduced in Group 1 variant mESC lines, and remained unchanged in Group 2 variant mESC lines (**Figure 3C**). Phosphorylation of AKT (Thr308) was reduced in all Group 1 variant mESC lines, but none of the Group 2 ones, consistent with our previous work (Baghdadi et al. 2025) (**Figure 3C**). These results indicate a reduction in mTORC1 activity in all variant mESC lines following stimulation of the IIS/mTOR signalling pathway, with a potential Group 1-specific PI3K dependent effect on AKT (Thr308). In addition, mTORC2 activity shows a Group 1-specific downregulation.

In the MAPK/ERK pathway, extracellular stimuli trigger the activation of upstream RAS proteins, which is followed by a phosphorylation cascade involving RAF proteins, MEK1/2, and ERK1/2 (Lavoie et al. 2020). In this study, we assessed MAPK/ERK pathway activity by MEK1/2 (Ser217/Ser221) and ERK1/2 (Thr202/Tyr204) phosphorylation in the normal state. Phosphorylation of both MEK1/2 (Ser217/Ser221) and ERK1/2 (Thr202/Tyr204) was reduced in all variant mESC lines, compared to the wild-type (**Figure 3D**). A comparable reduction was observed after insulin stimulation (**Figure S2**). These findings are consistent with the observations that we previously made for the MAPK/ERK pathway variants in the normal state and following insulin or EGF stimulation (Baghdadi et al. 2025) and indicate a reduction in MAPK/ERK signalling across all variant mESC lines, both in the normal state and following specific stimulation.

### Variants show distinct patterns downstream of MAPK/ERK signalling

In order to determine the effects of the identified variants downstream of the IIS/mTOR and MAPK/ERK signalling pathways, we assessed the expression of the ETS transcription factor family members and *Foxo3* in our transcriptomics dataset (**Figure 4A**). The ETS transcription factor family has been demonstrated to play a pivotal role in the regulation of lifespan in worms and flies (Dobson et al. 2019; Alic et al. 2014) and is regulated by the MAPK/ERK pathway (Wasylyk et al. 1998). In flies, MAPK/ERK signalling pathway activity has been shown to result in the activation of the transcriptional activator *Pnt*. The closest mammalian orthologues to *Pnt* are *Ets1* and *Ets2* (Vivekanand 2018). *Etv6*/*Tel* is the mammalian orthologue of transcriptional repressor *Aop*/*Yan* (Roukens et al. 2008). In flies, lifespan extension through a downregulation of RAS or ERK activity has been found to be Aop-dependent (Slack et al. 2015). Our results demonstrated that *Ets1* was upregulated in Group 1 and downregulated in Group 2 variant mESC lines, in comparison to the wild-type mESCs. Conversely, *Ets2* was upregulated in Group 2 variant mESC lines. qPCR analysis for *Ets1* and *Ets2* corroborated these observations in the set of five variant mESC lines (**Figure S4**), and was consistent with our previous work (Baghdadi et al. 2025). *Etv1* was downregulated in both variant groups, while *Etv4* was downregulated exclusively in Group 2, *Etv5* and *Etv6* did not demonstrate any significant deregulation in transcriptomics. These findings confirm a deregulation of the ETS transcription factor family, with both shared and distinct effects between the two groups of variants. In addition, together with the effects observed on mTORC1, mTORC2, MEK, and ERK activity, this indicates that variants located in genes involved in the IIS/mTOR signalling pathway can influence the activity of the MAPK/ERK signalling pathway activity, and vice versa.

**Figure 4.**
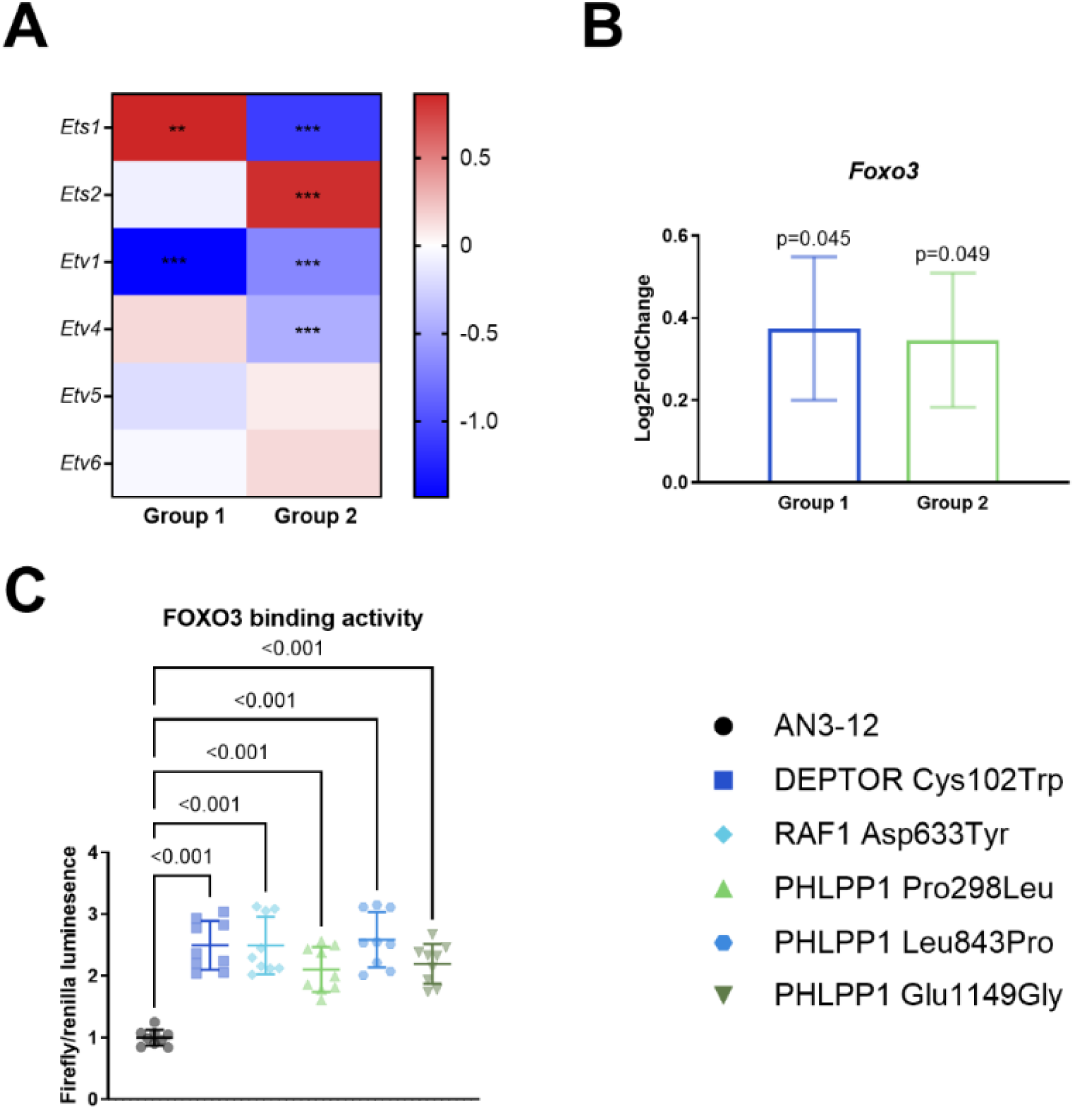
Variants alter the expression of the ETS transcription factors and *Foxo3*, and FOXO3 binding activity. Pathway activity assessments in transcriptomics (**A, B**) and transcription factor binding assay (**C**). **A** Transcript levels of ETS transcription factors (i.e., *Ets1*, *Ets2, Etv1, Etv4, Etv5, and Etv6*) in variant groups. *Ets1 and Ets2* show opposing deregulations between the two groups of variant mESC lines (Group 1: DEPTOR^Cys102Trp^, RAF1^Asp633Tyr^, and PHLPP1^Leu843Pro^; Group 2: IRS1^Pro4Thr^, IRS1^Tyr465His^, NF1^Phe1110Leu^, PHLPP1^Pro298Leu^, and PHLPP1^Glu1149Gly^). *Etv1* is downregulated in both variant groups. *Etv4* is downregulated only in Group 2. *Etv5* and *Etv6* show no significant deregulation. Change in expression compared to the wild-type is shown as Log2 fold change. **B** Increase in *Foxo3* in both variant groups. Change in expression compared to the wild-type is shown as Log2 fold change and error bars represent the standard error. **C** Increase in FOXO3 binding activity in all variant mESC lines. FOXO3 binding activity assessment through vector with firefly luciferase under the control of FHRE binding element (FHRE-luc). Renilla luciferase plasmid (pGL4.75) as control for transfection efficiency and cell number. In panel **C**, dots represent technical replicates, while error bars represent standard deviations. Data were analysed using a one-way ANOVA and Dunnett’s post hoc test.

### Consistent upregulation of FOXO3 activity in variant mESC lines

The interaction of the IIS/mTOR and MAPK/ERK signalling pathways with FOXO3, which has been identified to be important in the regulation of lifespan in humans (Willcox et al. 2008; Flachsbart et al. 2009; Flachsbart et al. 2017) and model organisms (Kenyon et al. 1993; Alic et al. 2014), motivated us to assess *Foxo3* transcript levels. Our results demonstrate that *Foxo3* levels were upregulated in all variant mESC lines compared to wild-type mESCs (**Figure 4B**). Given that transcript levels are an indirect measure of FOXO3 activity, we additionally measured FOXO3 binding activity using the FHRE-luc plasmid. The plasmid contains a luciferase reporter construct under the control of a basal promoter with three FHRE-binding elements (Brunet et al. 1999). All variant mESC lines exhibited increased FOXO3 binding activity in comparison to the wild-type mESCs, indicating enhanced FOXO3 activity (**Figure 4C**).

### Variant groups show opposing effects on growth signalling

Given the importance of the IIS/mTOR and MAPK/ERK signalling pathways in regulating cell growth and proliferation (Zhang and Liu 2002; Laplante and Sabatini 2012), we used the Incucyte to measure confluence over time. The confluence of Group 1 variant mESC lines increased less over time in comparison to wild-type, whereas Group 2 variant mESC lines exhibited the opposite (**Figure 5A**). We observed a corresponding pattern in our previous work (Baghdadi et al. 2025), indicating that the Group 1 variants cause a decrease in growth rate, while Group 2 variants increase it.

**Figure 5.**
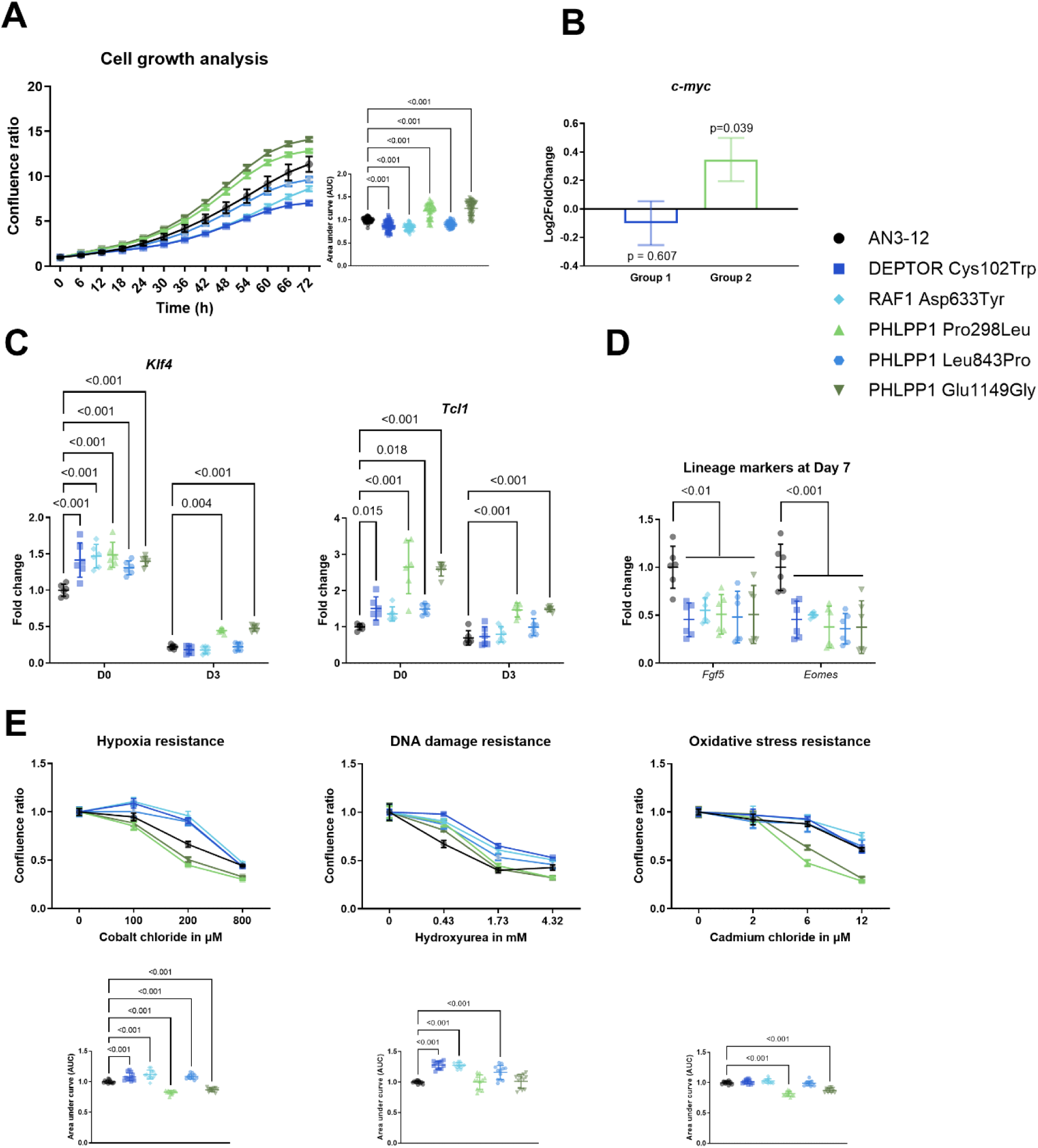
Variants affect growth pathway activity, pluripotency, differentiation potential, and resistance to stressors. **A** Cell growth over 72 hrs was assess by confluence and normalised to time point zero. Group 1 variants (DEPTOR^Cys102Trp^, RAF1^Asp633Tyr^, and PHLPP1^Leu843Pro^) showed slower confluence increases; Group 2 variants (PHLPP1^Pro298Leu^ and PHLPP1^Glu1149Gly^) showed the opposite. Left panel: one experiment (12 technical replicates). Right panel: pooled AUC from six independent experiments. **B** Increased *c-myc* transcript levels in Group 2 mESC variant lines in transcriptomics. Expression change compared to the wild-type is shown as Log2 fold change. **C** qPCR of pluripotency markers (*Klf4*, *Tcl1*) under normal growth and during spontaneous differentiation (LIF withdrawal). Normalised to wild-type at Day 0. Variant lines showed elevated pluripotency markers in normal conditions. After 3 days without LIF, Group 2 variants maintained higher levels. **D** qPCR of lineage markers (*Fgf5* – neuroectoderm; *Eomes* – endoderm) after 7 days without LIF. Normalised to wild-type under same conditions. All variants showed reduced lineage marker expression. *Rpl37a* was used for normalisation of all stemness qPCRs. Two independent experiments with 3 technical replicates each are shown. **E** Measurement of resistance to hypoxic, DNA damage, and oxidative stress. Confluence was measured 22 hrs after addition of stressor and normalised to un-stressed controls. Group 1 variants showed enhanced resistance to hypoxic and DNA damage stress. Group 2 variants showed a reduced resistance to hypoxic and oxidative stress. Upper panel: one experiment with 4 technical replicates per condition. Lower panel: pooled AUC from three independent experiments. For scatter plots, dots represent technical replicates. For growth curves and stress response curves, dots represent the mean of the technical replicates. Error bars represent standard deviations. Data were analysed using a one-way ANOVA (**A**, **B**, **D**, **E** – lower panels) or a two-way ANOVA (**C**) and Dunnett’s post hoc test. \**P* < 0.05, \*\**P* < 0.01, \*\*\**P* < 0.001.

Based on the observed differences in growth, we assessed *c-myc* transcript levels in our transcriptomics dataset (**Figure 5B**). In accordance with our previous work (Baghdadi et al. 2025), and the validation using qPCR (**Figure S4**), *c-myc* was specifically upregulated in Group 2 variant mESC lines. This finding indicates an increase in RNA polymerase transcription, ribosomal biogenesis, and cell growth (Poortinga et al. 2004).

### Variant mESC lines show a signature of increased pluripotency

Given that we selected a stem cell line as our model, we were interested in examining if the variants affect the cell’s pluripotency and differentiation potential (**Figure 5C-D**). Spontaneous differentiation was initiated by the withdrawal of LIF, based on a previously established protocol for the AN3-12 line (Kroef et al. 2025). In complete growth medium all variant mESC lines exhibited increased levels of the pluripotency markers *Klf4* and *Tcl1* (**Figure 5C**). After three days in -LIF medium, only Group 2 variant mESC lines showed elevated *Klf4* and *Tcl1* transcript levels. Lineage markers were reduced in all variant mESC lines following seven days of exposure to -LIF medium (**Figure 5D**). An experiment with only wild-type PHLPP1^Leu843Pro^ and PHLPP1^Glu1149Gly^ mESC lines and measurements of pluripotency markers at day 0, 1, and 3, as well as lineage markers at day 0, 3, and 7 exhibited the same patterns (**Figure S5**). The data thus suggests that the variant mESC lines exhibited an increased pluripotency in their normal state and differentiate less or slower after LIF withdrawal.

### Group 1 variants have an increased resistance to hypoxic and DNA damage stress

A number of models of life- and healthspan increasing interventions associated with the IIS/mTOR and MAPK/ERK signalling pathways demonstrated increased stress resistance to various stressors (Leiser et al. 2006; Murakami et al. 2003; Salmon et al. 2005; Johnson et al. 2002; Murakami 2006; Lithgow and Walker 2002). Utilising confluence as a viability metric in stress resistance assays, we observed group-specific effects on the response to different stressors (**Figure 5E**). Group 1 variant mESC lines showed an increased survival following exposure to hypoxic (cobalt chloride) and DNA damage (hydroxyurea) stress, while Group 2 variants exhibited reduced survival in response to exposure to hypoxic and oxidative (cadmium chloride) stress. Furthermore, we observed an inverse correlation between cell death and the changes in confluence for the majority of the conditions (**Figure S6**). In accordance with our previous work (Baghdadi et al. 2025), these observations confirm that Group 1 variants cause elevated stress resistance, a phenomenon not observed in Group 2 variants, which, dependent on the stressor, even showed an opposite effect.

### The variant mESC lines exhibit changes in lipid class abundance, and in mitochondrial metabolism and abundance

The IIS/mTOR and MAPK/ERK signalling pathways are growth pathways that regulate anabolism and proliferation (Burchfield et al. 2025). Growth is coupled to the availability of metabolites, other building blocks such as proteins, and energy. Cancer studies have shown that changes in IIS/mTOR genes and growth genes like *c-myc* cause extensive metabolic rewiring of the cells (Hsieh et al. 2015). Thus, we investigated the variants’ effects on metabolic regulation. We assessed lipid and mitochondrial metabolism as ORA on transcriptomics and proteomics indicated their deregulation.

Motivated by the enrichment of the hallmark “Cholesterol homeostasis” in the functional enrichment analysis, we performed lipidomics on the same 10 variant mESC lines as for the previous omics (**Figure 1** and **Table 2**). The PCA of the lipidomics data demonstrated the same grouping of the variants into two distinct groups (**Figure 6A**). A comparison of abundance of lipid classes per group against the wild-type revealed that the majority of changes in lipid classes were consistent between both groups.

**Figure 6.**
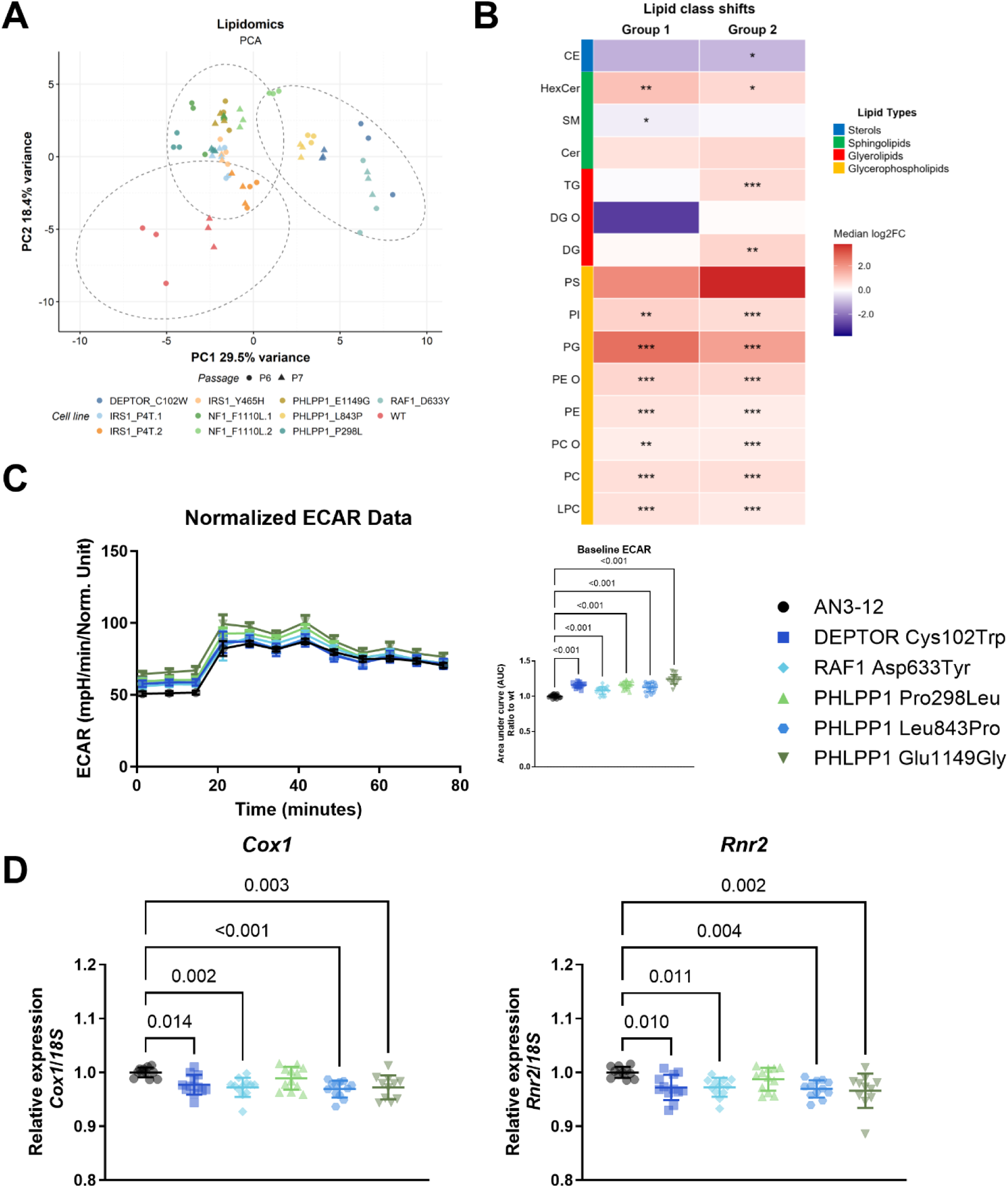
Variants affect lipid and mitochondrial metabolism, and mitochondrial abundance. **A**, **B** Lipidomic analysis of the extended set of variant mESC lines. Variants cluster into two groups both distinct from the wild-type in PCA (**A**). **B** Changes in lipid class in Group 1 and Group 2 of variant mESC lines (Group 1: DEPTOR^Cys102Trp^, RAF1^Asp633Tyr^ and PHLPP1^Leu843Pro^; Group 2: IRS1^Pro4Thr^, IRS1^Tyr465His^, NF1^Phe1110Leu,^ PHLPP1^Pro298Leu^ and PHLPP1^Glu1149Gly^). Betas represent the Log2 fold change compared to the wild-type. **C** Extracellular acidification rate (ECAR) was measured in Seahorse in a Mito Stress Test-assay. The left panel shows one experiment with 10 technical replicates. The right panel shows a combination of AUC analysis of baseline ECAR (measurement 1-3) of two independent experiments. Baseline ECAR was increased in all variant mESC lines. **D** mtDNA copy number analysis via qPCR on total DNA. Mitochondrial probes (*Cox1* and *Rnr2*) were normalised to genomic DNA copy number (*18S*). All variant mESC lines, except PHLPP1^Pro298Leu^, show a reduction in mtDNA copy number. For scatter plots, dots represent technical replicates. For the normalised ECAR curve, dots represent the mean of the technical replicates. Error bars represent standard deviations. Data were analysed using a one-way ANOVA and Dunnett’s post hoc test. \**P* < 0.05, \*\**P* < 0.01, \*\*\**P* < 0.001. *CE*; Cholesteryl Esters, *Cer*; Ceramides, *SM*; Sphingomyelins, *HexCer*; Hexosylceramides, *DG*; Diacylglycerols, *DG.O*; Ether Diacylglycerols, *TG*; Triacylglycerols, *LPC*; Lysoglycerophosphatidylcholines, *PC*; Glycerophosphatidylcholines, *PC.O*; Ether Glycerophosphatidylcholines, *PE*; Glycerophosphatidylethanolamines, *PE.O*; Ether Glycerophosphatidylethanolamines, *PG*; Glycerophosphatidylglycerols, *PI*; Glycerophosphatidylinositols, *PS*; Glycerophosphatidylserines.

Glycerophospholipids were consistently upregulated in both groups and all detected lipid classes. Additionally, Hexosylceramides (HexCers) were upregulated and Cholesterol Esters (CEs) showed a (trend of) downregulation in both variant groups. Moreover, Triacylglycerols (TGs) and Diacylglycerols (DGs) were specifically upregulated in Group 2 variant mESC lines, while Sphingomyelins (SMs) show a specific downregulation in Group 1 variant mESCs (**Figure 6B**). These findings demonstrate changes in various aspects of lipid metabolism for all variant mESC lines.

The enrichment of the hallmark “Glycolysis” motivated us to investigate mitochondrial metabolism and abundance. With the Seahorse Mito Stress Test-assay we could only detect a mild decrease in the baseline oxygen consumption rate (OCR), which is a measure for the activity of mitochondrial oxidative phosphorylation (Plitzko and Loesgen 2018), of the PHLPP1^Glu1149Gly^ variant mESC line (**Figure S7**). However, the baseline extracellular acidification rate (ECAR), which reflects glycolytic activity (Plitzko and Loesgen 2018), was elevated in all variant mESC lines (**Figure 6C**). Furthermore, mtDNA copy numbers were significantly reduced in all variant mESC lines with the exception of the PHLPP1^Pro298Leu^ variant mESC line (**Figure 6D**). These findings indicate mild effects on mitochondrial metabolism for all variant mESC lines.

## Discussion

In this study, we showed that rare genetic variants in the IIS/mTOR and MAPK/ERK signalling pathways identified in exceptionally long-lived individuals show shared and distinct *in vitro* effects in assays representing longevity characteristics in cellular and animal models. The variants clustered in two distinct groups apart from the wild-type along the axes representing the largest variances (PC1 and PC2) in a PCA for both transcriptomic and proteomic datasets. All variant mESC lines exhibited a reduction in mTORC1 and MAPK/ERK signalling pathway activity. An increase in *Foxo3* expression and FOXO3 activity, as well as an increase in ECAR, were also shared between all variant mESC lines. The majority of variants displayed a reduction in mtDNA levels. The alterations in the abundance of lipid classes were mostly shared between both groups of variants. Pluripotency markers were increased, while differentiation markers were reduced after inducing spontaneous differentiation in all variant cell lines. For stress resistance and growth speed, variant groups behaved differently. Group 1 showed a reduction in growth speed and an increased resistance to some stressors compared to the wild-type. Conversely, Group 2 variants exhibited an increased growth speed and reduced resistance to certain stressors.

We identified functional effects that are common between all transgenic mESC lines carrying variants identified in two different studies. It is notable that several of these common effects can be linked to known mechanisms involved in the regulation of lifespan in animal models. This aligns with the enrichment of rare variants identified in Ashkenazi Jewish centenarians within the insulin signalling network (Lin et al. 2021), although functional readouts were not assessed in that study. One of the common effects observed for all variant mESC lines is the reduction of IIS/mTOR and MAPK/ERK signalling pathway activity. The downregulation of these pathways has been observed to associate with mouse models exhibiting an increased life- and healthspan (Selman et al. 2009; Jiang et al. 2023; Sun et al. 2009; Wink et al. 2022). In our previous work, we demonstrated that variants in the MAPK/ERK pathway were associated with a downregulation of both the MAPK/ERK and IIS/mTOR signalling pathways (Baghdadi et al. 2025). In this study, we show that this cross-talk also occurs in the opposite direction. Variants located in both the IIS/mTOR and the MAPK/ERK signalling pathways downregulate the activity of both pathways and affect their downstream targets, the ETS transcription factor family and FOXO3. The final effector of the Ras/MEK/ERK cascade, ERK, functions as a negative regulator of FOXO3 (Tian et al. 2022; Yang et al. 2008), linking the observed downregulation of MAPK signalling and upregulation of *Foxo3* transcript levels and FOXO3 binding activity. The upregulation of FOXO3 activity appears to be inconsistent with the upregulation of AKT (Ser473) phosphorylation in the normal state, indicating increased mTORC2 activity. mTORC2 has been identified as a negative regulator of FOXO3 activity (Jimenez et al. 2022). However, it can be speculated that the upregulation of AKT (Ser473) phosphorylation is due to the relief of a negative feedback and upregulation of upstream PI3K signalling that has been observed when mTORC1 is inhibited through, for example, rapamycin (Sun et al. 2005; O’Reilly et al. 2006; Shi et al. 2005). FOXO3 has been shown to positively associate with longevity and healthy ageing across organisms (McIntyre et al. 2022; Kenyon et al. 1993; Inci et al. 2025; Willcox et al. 2008). In worms, the lifespan-extending effect of the downregulation of IIS/mTOR signalling, through daf-2 (worm orthologue of IGF1R), was found to be dependent on daf-16 (worm orthologue of FOXOs) (Kenyon et al. 1993). Furthermore, FOXO3 single-nucleotide polymorphisms (SNPs) that were associated with longevity in three European populations (including the GLS) demonstrated enhanced promoter activity *in vitro* as well as an increase of *Foxo3* mRNA expression in the blood of the carriers (Flachsbart et al. 2017). Further investigations are required to determine whether the other effects observed here are also dependent on FOXO3.

The elevated pluripotency signature that we observed for all variants can be attributed to the significance of the IIS/mTOR and MAPK/ERK signalling pathways in the regulation of pluripotency. Inhibition of MAPK/ERK signalling has been demonstrated to be associated with increased stemness (Hishida et al. 2015), while the activity of p70 S6K has previously been linked to reduced pluripotency and increased differentiation (Romanyuk et al. 2024; Kim et al. 2024). However, it should be noted that this phenotype is likely to be specific to stem cells and not transferrable to other cell types.

We used untargeted proteomics and transcriptomics in order to identify the mechanisms affected by the variants. Among others, functional follow up showed effects on lipid and mitochondrial metabolism. Both an increase in baseline ECAR and the reduction in mtDNA levels can be linked to an increased glycolytic rate (Yoo et al. 2024; Mou et al. 2018) and may thus explain the enrichment of the hallmark “Glycolysis” in both the proteomics and transcriptomics analyses. An increase in glycolytic rate would be concomitant with the increased expression of *Klf4* and *Tcl1*, as both of these are crucial in directing stem cell metabolism towards glycolysis (Nishimura et al. 2017). However, the effects on mitochondrial metabolism observed here do not seem to be stem cell-specific. mtDNA measurements in blood samples from nonagenarian siblings from the LLS, their middle-aged offspring, and control individuals, revealed a reduction in mtDNA copy number for both the exceptionally long-lived individuals and their offspring (van Leeuwen et al. 2014). Thus, the reduction of mtDNA levels in the variant mESC lines aligns with the hypothesis that lifespan-increasing interventions and genetic variants can act through a preservation of mitochondrial function rather than enhanced mitochondrial biogenesis (Lanza et al. 2012; van Leeuwen et al. 2014).

The effects in lipid metabolism in the variant mESCs is consistent with the previously identified importance of lipid metabolism in the regulation of lifespan in several model organism studies (Johnson and Stolzing 2019). For example, the metabolite group that was affected most by the lifespan-increasing drugs rapamycin, acarbose, 17α-estradiol, and canagliflozin in mice was lipids (Greenfield et al. 2025). These drugs are known to attenuate the age-related increase in MAPK/ERK signalling in mice (Wink et al. 2022; Jiang et al. 2023). These findings establish a link between their effects on lifespan and their impact on both lipid metabolism and MAPK/ERK signalling, effects that were also observed in the variant mESC lines. Studies on the middle-aged offspring of exceptionally long-lived individuals in the LLS have implicated lipid metabolism as an important factor for familial longevity (Vaarhorst et al. 2011; Gonzalez-Covarrubias et al. 2013). Additionally, genetic variants associated with cholesterol metabolism have been identified as a potential key for a healthy ageing trajectory in families from the LLS (Sant’Anna Barbosa Ferreira et al. 2025), which would be in line with the enrichment of the hallmark “Cholesterol Homeostasis” in both groups in the transcriptomics data analysis. The changes in lipid species we observed in the current study add to the previously identified importance of lipid metabolism in human familial longevity.

In addition to changes in lipid metabolism and a decreased mtDNA copy number, a favourable glucose metabolism was identified as a signature of familial longevity in the LLS (Rozing et al. 2010; Wijsman et al. 2011). The changes in IIS/mTOR signalling and FOXO3 activity that we observed here can be linked to effects of beneficial glucose metabolism observed in long-lived mice and humans (Banasik et al. 2011; Selman et al. 2009). Thus, we can draw clear connections between the metabolic signatures of familial longevity in the LLS and the effects that we observe in the variant mESC lines.

For both proteomics and transcriptomics, the variants cluster into two groups based on PC1, which are both different from the wild-type cells based on PC2. A similar PCA grouping can be observed for the lipidomics. Interestingly, this clustering cannot be explained by the pathway or the gene in which the variants are residing, nor the study in which they were identified. Despite our main interest in the shared effects between all variants, it is interesting to characterise which effects are distinct between the two variant groups. Both variant groups may elicit shared beneficial effects, as well as additional group-specific beneficial effects. A limitation of this study is, however, that we do not yet understand the reason for the observed grouping. Characterisation of the variants in an additional cell line could elucidate if this grouping is a stem cell-specific effect. One of the functional readouts that shows differential effects between variant groups is resistance to stressors. The elevated resistance towards hypoxia and DNA damage stress for Group 1 variants is in line with increased stress resistance observed in fibroblasts isolated from long-lived pituitary dwarf, Snell dwarf and Ames dwarf mice (Leiser et al. 2006; Murakami et al. 2003; Salmon et al. 2005). However, there are also reports of lifespan-increasing cellular and worm models that do not cause changes or even a reduction in resistance to some stressors (Page et al. 2014; Soo et al. 2023; van Raamsdonk and Hekimi 2009). This finding supports the hypothesis that lifespan and stress resistance are correlated, yet not invariably physiologically related (Amrit et al. 2019; Dues et al. 2019). From these findings we can conclude that the reduced resistance to hypoxia and oxidative stress in Group 2 variant mESC lines does not contradict the potential contribution of these variants to the longevity of their carriers.

Another functional readout on which we observed divergent effects is growth speed. A signature associated with an increased growth rate, exclusive to Group 2, is the increase in *c-myc* transcripts. MYC is a downstream effector of the MAPK/ERK signalling pathway (Zuo et al. 2023) and known for its role in the regulation of RNA polymerase transcription, ribosomal biogenesis, and cell growth (Poortinga et al. 2004). Thus, the upregulation in Group 2 is in line with the increased growth rate in Group 2 variant mESCs, but unexpected given the downregulation of MAPK/ERK signalling. In light of MYC’s function in ESC pluripotency and self-renewal (Cartwright et al. 2005; Varlakhanova et al. 2010), it could be hypothesised that the effect on *c-myc* transcripts and growth speed in Group 2 variants is stem cell-specific. Following three days of LIF withdrawal, Group 2 variants, unlike Group 1 variants, exhibited an increase in *Klf4* and *Tcl1* transcripts. Together, this indicates that Group 2 variants might cause a partially LIF-independent effect that leads to increased pluripotency, even in the absence of LIF. The elevated growth rate observed in Group 2 variants may signify an increased capacity of self-renewal.

The mESC line AN3-12, which we selected as a model here, is particularly suitable for CRISPR/Cas9 gene editing, since it is a haploid cell line that spontaneously becomes diploid over time. However, this also resulted in one of the major limitations of this work. We functionally characterised the variants exclusively in a homozygous state, despite being identified as heterozygous in the exceptionally long-lived individuals. In future research, the impact of the variants in a heterozygous state should also be investigated. Furthermore, the characterisation of the variants in a somatic cell line would facilitate the comprehension of which of the effects identified here might be stem cell-specific. Ideally, this cell line would be of human origin, thus enabling the examination of the effects of the variants in a human context. In this study, only a single colony was successfully generated for the majority of the variants, which presents another limitation. However, the fact that the variants cluster together in two groups in the PCA and as one group for the majority of the functional readouts, makes it rather unlikely that the effects that we observe here are due to CRISPR off-target effects. The CRISPR protocol that we utilised here exhibited a certain degree of inefficiency. However, this was counterbalanced by the use of the precise nickase mutant Cas9n. In the event of similar experiments in the future, advancements in the CRISPR/Cas technology, such as the use of non-homologous end joining (NHEJ) inhibitors, have the potential of enhancing efficiency without reducing specificity. In order to strengthen the *in vitro* characterisation performed here, *in vivo* characterisation in mice, such as lifespan and healthspan assessment or insulin sensitivity measurements could be performed to complement our results. It would also be ideal to conduct follow-up measurements on primary cells from the exceptionally long-lived individuals, but these are not available for either the LLS or the GLS. A further consideration for subsequent experiments should be the utilisation of phosphoproteomics. In this study, the assessment of pathway activity through phosphorylation of a protein was limited to a specific set of targeted proteins using western blotting. Phosphoproteomics has the potential to provide a more comprehensive and untargeted assessment of pathway activity. This approach may facilitate the identification of changes in pathway activity that could potentially underlie the observed phenotypes. Moreover, in future studies, a hypothesis-free approach may be employed to identify novel players and pathways in the regulation of longevity. A recent example of this is the identification and in vitro characterisation of a cGAS variant identified via sib-pair linkage analysis in the LLS (Putter et al. 2025).

## Conclusions

We employed a candidate pathway approach to study the *in vitro* functional effects of a selection of rare variants in the IIS/mTOR and MAPK/ERK signalling pathways carried by exceptionally long-lived individuals. In conjunction with our preceding publication (Baghdadi et al. 2025), we have demonstrated that the mESC line AN3-12 can serve as an effective model for the investigation of rare genetic variants *in vitro*. With the present study we demonstrate the functional interplay between the IIS/mTOR and MAPK/ERK signalling pathways through human genetic variants, which could explain why targeting of each of the two pathways in model organisms leads to similar lifespan-related effects. We showed that the variants identified in exceptionally long-lived individuals have functional effects that are analogous to those observed in animal models with increased lifespan and improved healthspan. Additionally, we can draw connections between metabolic signatures of familial longevity observed in the LLS with some of the observations we made in the variant mESC lines. Subsequent analyses of the shared effects of the variants in *in vivo* models can be used to determine whether these individual variants are sufficient to extend lifespan. In consideration of these findings, interventions could be designed to recapitulate the effects common to all variants and thereby mimic the effects underlying the improved life- and healthspan in exceptionally long-lived individuals.

## List of abbreviations

CADD: Combined Annotation Dependent Depletion
CE: Cholesteryl Esters
Cer.O2: Dihydroceramides
DG: Diacylglycerols
DG.O: Ether Diacylglycerols
ECAR: Extracellular acidification rate
ERK: Extracellular signal-regulated kinase
FACS: Fluorescence-activated cell sorting
FHRE: Forkhead responsive element
GLS: German Longevity Study
gRNA: Guide RNA
GWAS: Genome-wide association studies
HexCer.O2: Hydroxylated Hexosylceramides
IIS: Insulin/insulin-like growth factor 1 signalling
LIF: Leukaemia inhibitory factor
-LIF medium: Medium without LIF
LLS: Leiden Longevity Study
LPC: Lysoglycerophosphatidylcholines
MAF: Minor allele frequency
MAPK: Mitogen-activated protein kinase
mESCs: Mouse embryonic stem cells
mtDNA: Mitochondrial DNA
MPI-Age: Max Planck Institute for Biology of Ageing
mTOR: Mechanistic target of rapamycin
NEHJ: Non-homologous end joining
OCR: Oxygen consumption rate
ORA: Overrepresentation analysis
PC: Glycerophosphatidylcholines
PC.O: Ether Glycerophosphatidylcholines
PCA: Principal component analysis
PE: Glycerophosphatidylethanolamines
PE.O: Ether Glycerophosphatidylethanolamines
PG: Glycerophosphatidylglycerols
PI: Glycerophosphatidylinositols
PS: Glycerophosphatidylserines
SM.O2: Hydroxylated Sphingomyelins
SNP: Single-nucleotide polymorphism
ssODN: Single-stranded DNA oligonucleotide
TG: Triacylglycerols
VST: Variance-stabilized transformed

## Declarations

### Availability of data and materials

The transcriptomics dataset generated and analysed during the current study are available in the NCBI Gene Expression Omnibus (GEO) repository under accession number GSE[Accession_Number]. The mass spectrometry proteomics data have been deposited to the ProteomeXchange Consortium via the PRIDE partner repository with the dataset identifier PXD078725, token: Xlr32OADI11U. The lipidomics data have been deposited to MetaboLights repository (Yurekten et al. 2024) with the study identifier MTBLS14592. Processed data are available upon reasonable request from the corresponding author Joris Deelen.

The customised script used for the analysis of the datasets in this article are available on GitHub (https://github.com/artonifil/ELLI_variants_in_mESCs.git).

Population allele frequency data were obtained from gnomAD v4.1 (GRCh38) (Chen et al. 2024) and assessed via the gnomAD browser (https://gnomad.broadinstitute.org/).

Variant deleteriousness was predicted using the CADD score v1.6 and v1.7 (Schubach et al. 2024) and assessed via the CADD browser (https://cadd.bihealth.org/).

## Contributions

Conceptualisation: MN, LS, PES, MBa, JD; Methodology: MN, LS, HH, RP, MBa, JD; Investigation: MN, LS; Formal analysis: MN, EB, DK, FA, IB, LB, HEDS; Visualisation: MN, FA, IB; Resources: AB, MBe, AN, PES, JD; Writing – original draft: MN; Writing – review and editing: MBa, JD; Supervision: AP, AB, AN, PES, MBa, JD; Funding acquisition: AB, AN, PES, JD. All authors read and approved the final manuscript.

## Acknowledgements

We thank the study participants of the Leiden Longevity Study and the German Longevity Study. We thank the Proteomics, FACS and Imaging, Bioinformatics, and Metabolomics Core Facilities of the MPE-Age for outstanding technical help and advice. We thank Virginia Kroef for the guidance for the AN3-12 cells and the CRISPR protocol. We thank Lea Hund for her guidance for the differentiation experiment. We thank Soni Deshwal for the guidance for the Seahorse XF Cell Mito Stress Test. We thank Elina Singer for the help in the preparation of the lipidomics samples. Figure 1 was created with BioRender.com.

