## Supplementary Information for "Rare genetic variants in the IIS/mTOR signalling pathway identified in exceptionally long-lived individuals show shared in vitro effects associated with lifespan across species"

Neuerburg, M. et al.

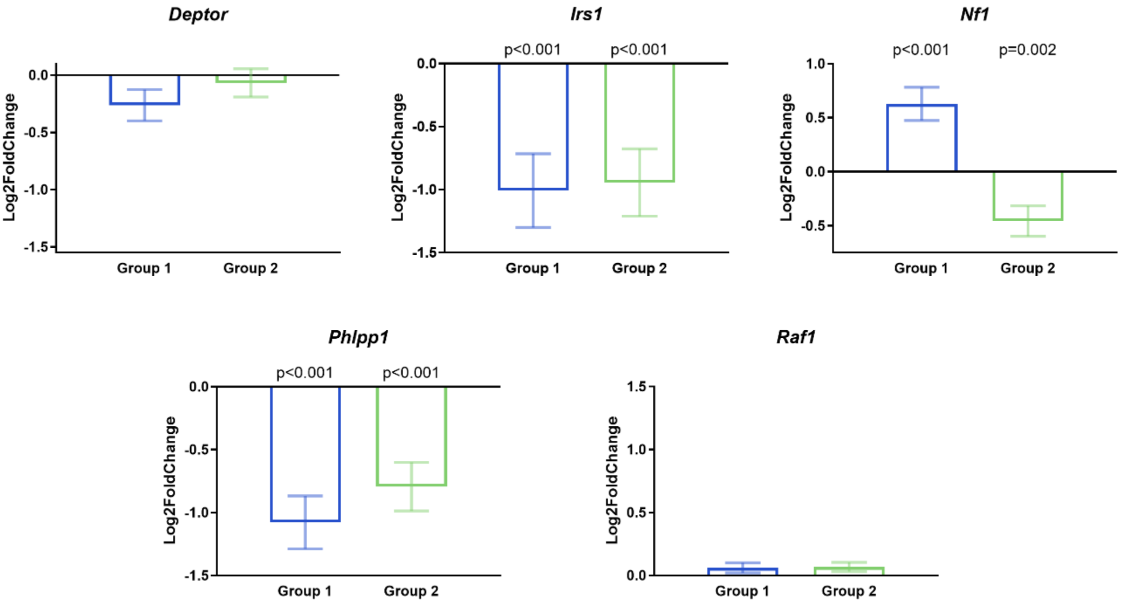


**Figure S1 | Expression of variant carrying genes in transcriptomics.** Decrease of *Phlpp1* and *Irs1* transcripts in both variant groups in transcriptomics. Upregulation of *Nf1* transcripts in Group 1 and downregulation in Group 2. No change in *Raf1* and *Deptor* transcripts in both groups. Change in expression compared to the wild-type is shown as Log2 fold change and error bars reflect the standard error.

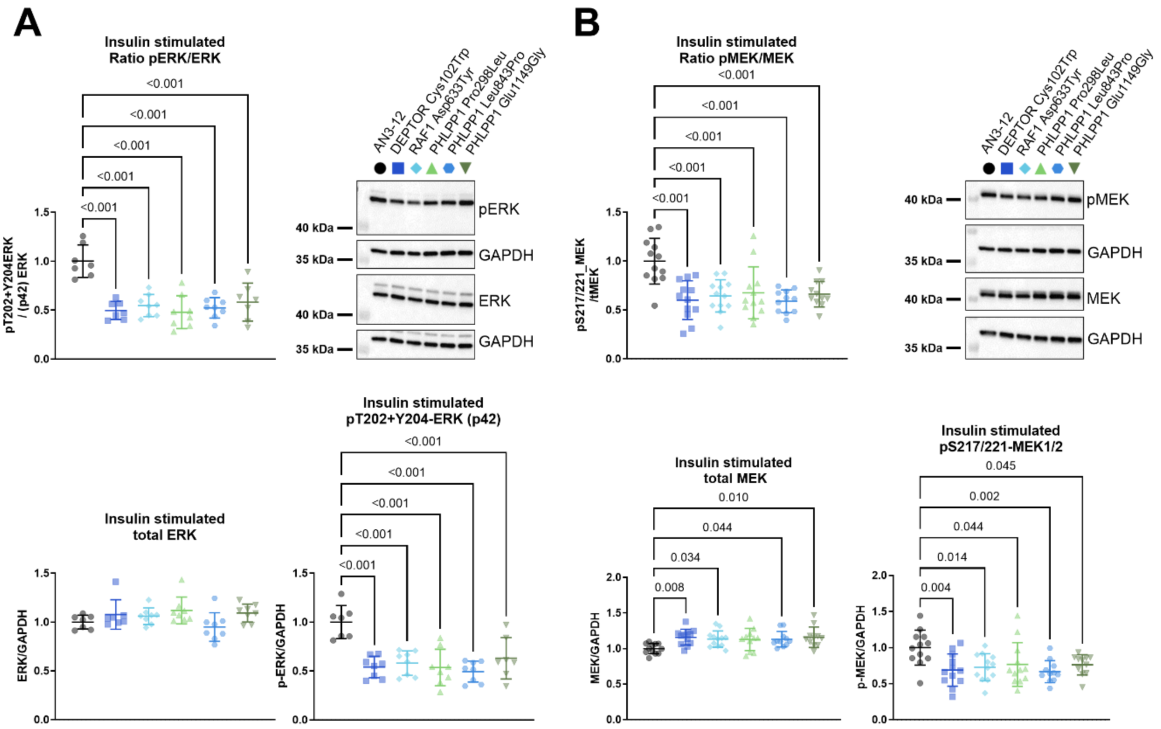


**Figure S2 | Reduced MAPK/ERK signalling after insulin stimulation.** Analysis of ERK (**A**) and MEK (**B**) phosphorylation ratios using western blotting. Upper panels show ratios of phosphorylated over total ERK and MEK respectively. mESCs were serum-starved for 6 hrs before a 10 min insulin stimulation (100 nM). Ratios, as well as phosphorylated protein for both ERK (**A**) and MEK (**B**) were downregulated. Total MEK is mildly increased for all variant mESC lines (**B**). For all experiments, GAPDH was used for normalisation. Data shown is from two to three independent experiments with four technical replicates each. Error bars represent standard deviations. Data were analysed using a one-way ANOVA and Dunnett's post hoc test.

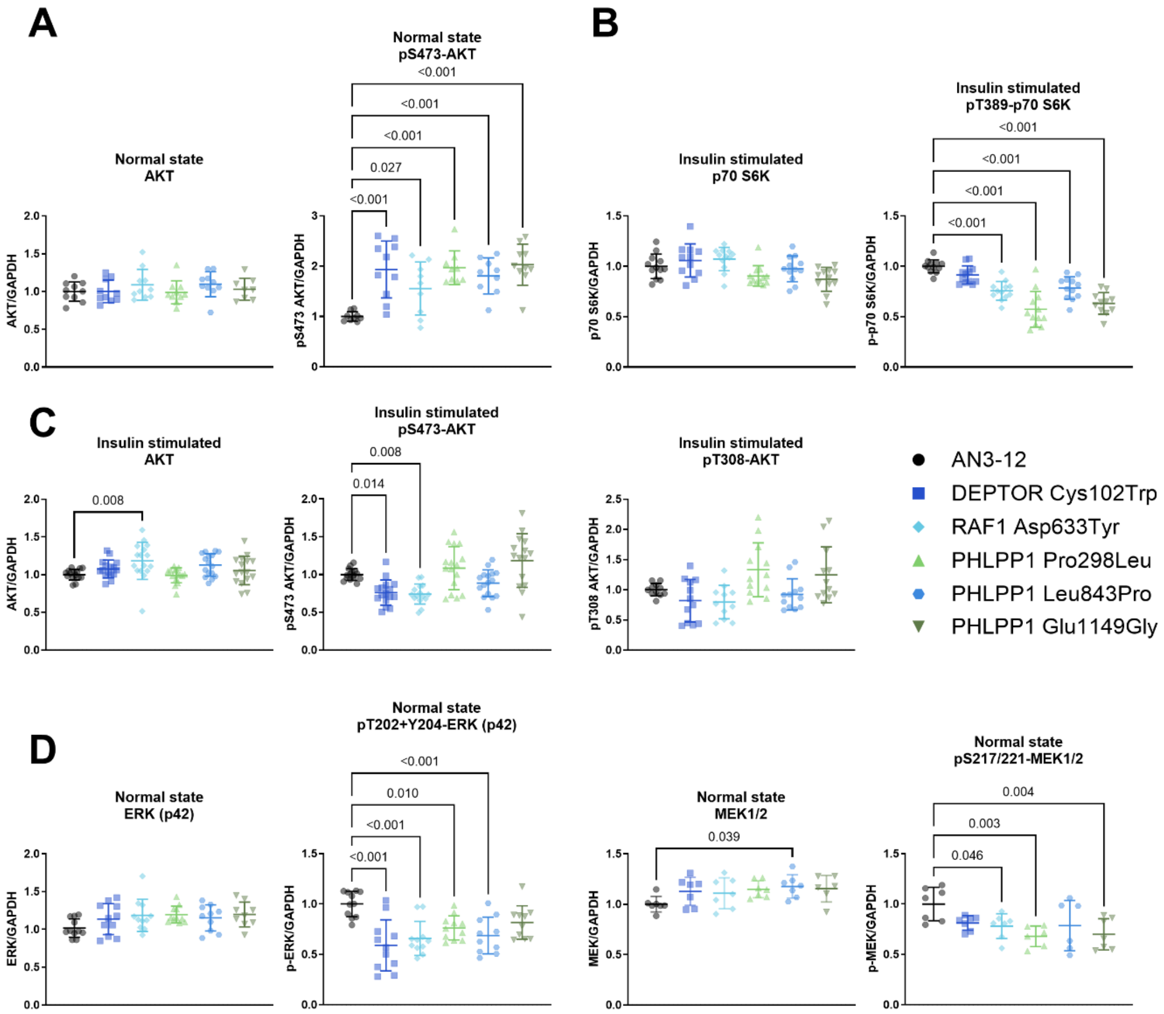


**Figure S3 | Quantification of total and phosphorylated proteins in IIS/mTOR and MAPK/ERK signalling pathways. A** Increase of phosphorylated AKT (Ser473) under normal growth conditions. **B** Decrease of phosphorylated p70 S6K (Thr389) after insulin stimulation. **C** Decrease of phosphorylated AKT (Ser473) in Group 1 variant mESC lines after insulin stimulation. Mild pattern of decrease for Group 1 and increase for Group 2 variants in phosphorylated AKT (Thr308) after insulin stimulation. Increase in total AKT in Group 1 variant mESC lines after insulin stimulation. **D** Decrease in phosphorylated ERK and MEK under normal growth conditions. Mild increase in total MEK in all variant mESC lines under normal growth conditions. Normal state phosphorylation was assessed in mESCs grown in complete growth medium (**A**, **D**). For insulin stimulation (**B**, **C**) mESCs were serum-starved for 6 hrs before a 10 min insulin stimulation (100 nM). For all experiments, GAPDH was used for normalisation. Data shown is from two to three independent experiments with four technical replicates each. Error bars represent standard deviations. Data were analysed using a one-way ANOVA and Dunnett's post hoc test.

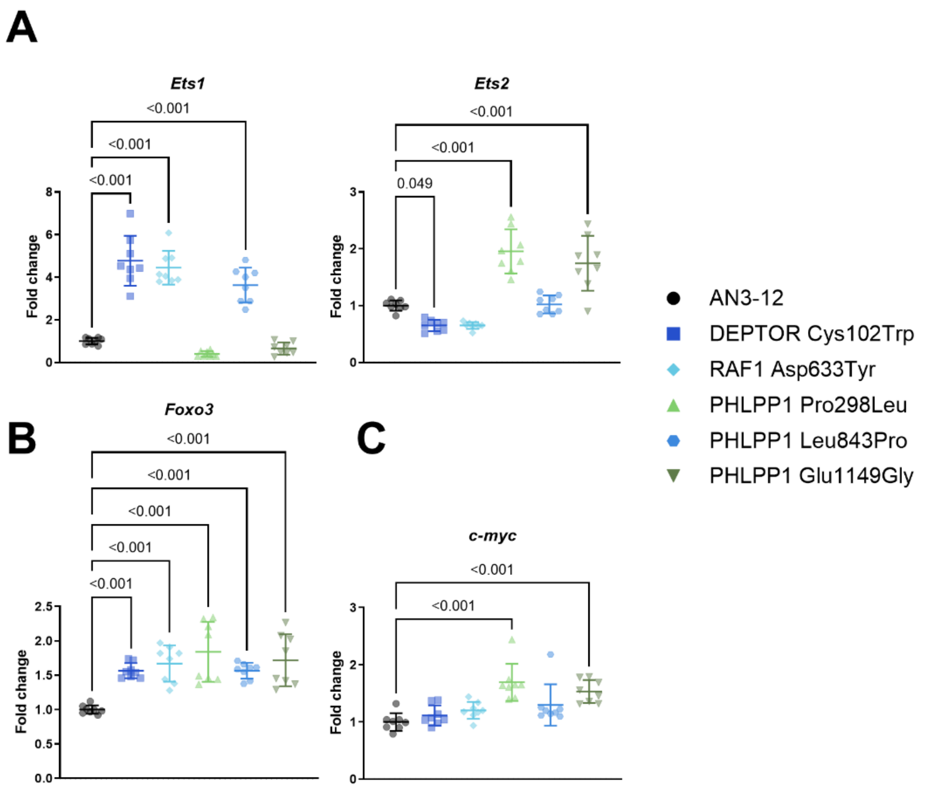


**Figure S4 | qPCR quantification of transcription factors.** qPCR of ETS transcription factors (i.e., *Ets1* and *Ets2*) (**A**), *Foxo3* (**B**), and *c-myc* (**C**) in variant mESC lines. **A** *Ets1* and *Ets2* show opposing deregulations between the two groups of variant mESC lines (Group 1: DEPTOR$^{Cys102Trp}$, RAF1$^{Asp633Tyr}$ and PHLPP1$^{Leu843Pro}$; Group 2: PHLPP1$^{Pro298Leu}$ and PHLPP1$^{Glu1149Gly}$). **B** Increase in *Foxo3* transcripts in all variant mESC lines. **C** Increase in *c-myc* transcript levels in Group 2 variant mESC lines in qPCR. *Gapdh* was used as a control. Two experiments with 3 technical replicates each, are shown. Dots represent technical replicates. Error bars represent standard deviations. Data were analysed using a one-way ANOVA and Dunnett's post hoc test.

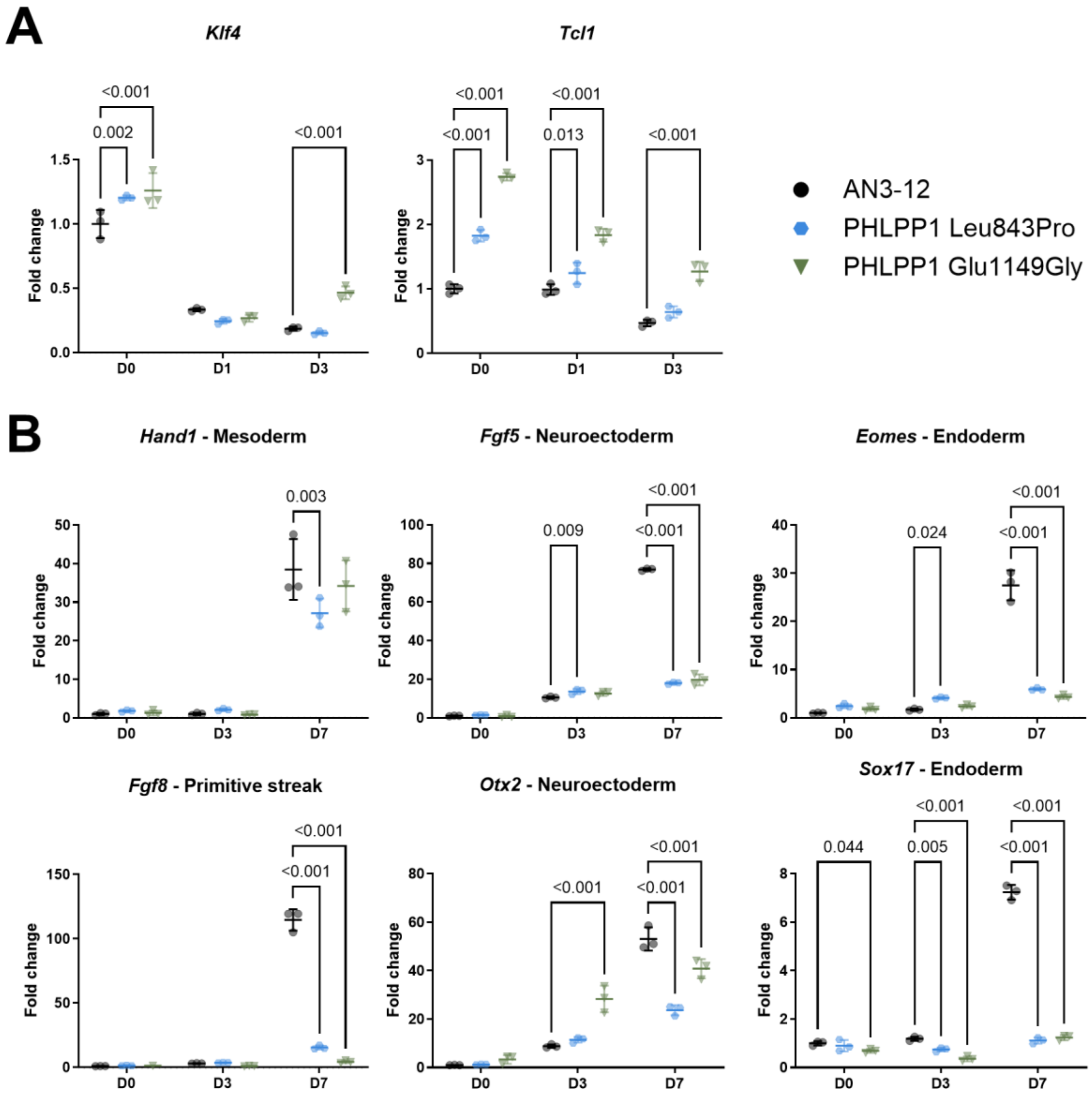


**Figure S5 | Measurement of pluripotency and differentiation potential.** qPCR analysis of pluripotency markers (*Klf4*, *Tcl1*) (**A**) and lineage markers (*Hand1* – mesoderm, *Fgf5* – neuroectoderm, *Eomes* – endoderm, *Fgf8* – primitive streak, *Otx2* – neuroectoderm, *Sox17* - endoderm) (**B**) on variant mESC lines after activation of spontaneous differentiation (i.e., withdrawal of LIF). All samples were normalised to the wild-type under normal growth conditions (Day 0). Pluripotency markers were measured before the withdrawal of LIF (D0) and one (D1) and three (D3) days after the withdrawal of LIF. Lineage markers were measured before the withdrawal of LIF (D0), three (D3) and seven (D7) days after the withdrawal of LIF. **A** LIF withdrawal caused a reduction in pluripotency markers for all three mESC lines. In normal growth conditions variant mESC lines showed increased pluripotency marker levels. One day after LIF withdrawal both variant mESC lines showed increased Tcl1 transcripts compared to the wild-type. At Day 3 the PHLPP1$^{Glu1149Gly}$ variant, mESC line showed increased pluripotency markers. **B** All mESC lines showed an increase in pluripotency marker expression after three to seven days of LIF withdrawal compared to the growth in normal growth medium. Seven days after LIF withdrawal both variant mESC lines showed a reduction in lineage markers compared to the wild-type. *Rpl37A* was used for normalisation. One experiment with three technical replicates is shown. Dots represent technical replicates. Error bars represent standard deviations. Data were analysed using a two-way ANOVA and Dunnett's post hoc test.

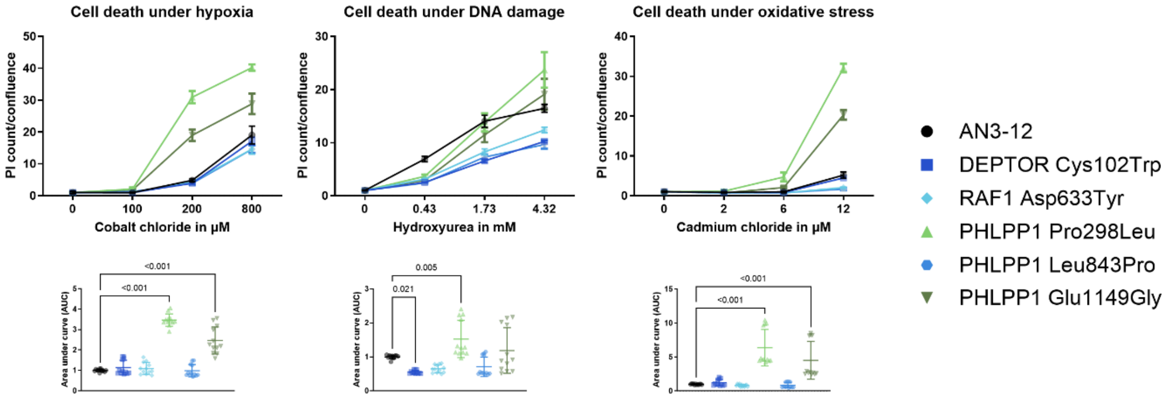


**Figure S6 | Cell death in stress resistance assays.** Cell death assessment under the exposure to hypoxic (cobalt chloride), DNA damage (hydroxyurea) and oxidative (cadmium chloride) stress. Confluence and propidium iodide count were measured 22 hrs after addition of stressor. A ratio of propidium iodide count over confluence was created and normalised to the control without drug of the respective cell line. Group 2 variants showed increased cell death under hypoxic and oxidative stress. Under DNA damage stress, Group 1 variant mESC lines showed reduced cell death while for Group 2 variants the results varied between replicates, with an average increase in cell death for the PHLPP1$^{Pro298Leu}$ variant mESC line. The upper panel shows one experiment with 4 technical replicates per condition. The lower panel shows the combination of the AUC measurement of three independent experiments. For scatter plots, dots represent technical replicates. For cell death curves, dots represent the mean of the technical replicates. Error bars represent standard deviations. Data were analysed using a one-way ANOVA (lower panels) and Dunnett's post hoc test.

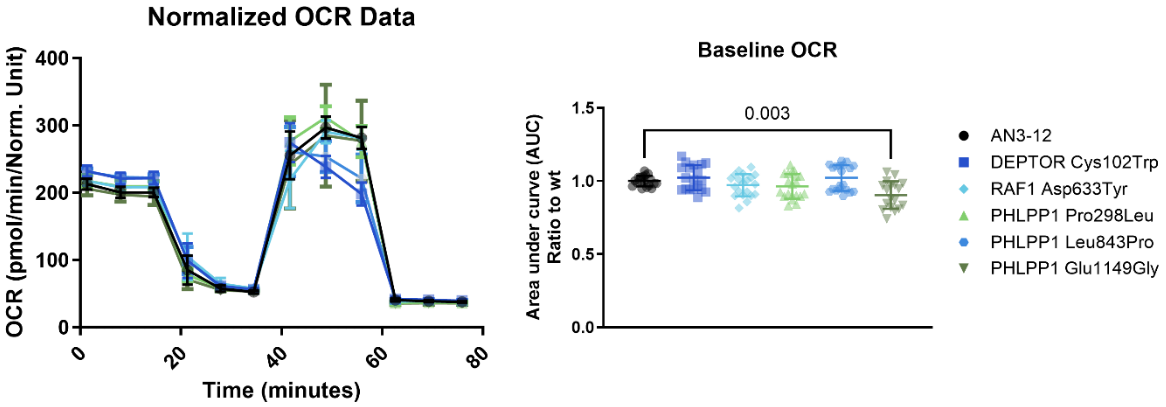


**Figure S7 | Measurement of the oxygen consumption rate.** Oxygen consumption rate (OCR) was measured in Seahorse in a Mito Stress Test-assay. The left panel shows one experiment with 10 technical replicates. The right panel shows a combination of AUC analysis of baseline OCR (measurement 1-3) of two independent experiments. Baseline OCR was mildly decreased in the PHLPP1$^{Glu1149Gly}$ variant and unchanged in all other variant mESC lines. For scatter plots, dots represent technical replicates. For the normalised OCR curve, dots represent the mean of the technical replicates. Error bars represent standard deviations. Data were analysed using a one-way ANOVA and Dunnett's post hoc test.

**Table S1 | Overview of gRNAs used for CRISPR of mESCs.**

| Target mutation | Vector backbone | gRNA1 | gRNA2 |
|---|---|---|---|
| DEPTOR Cys102Trp | pSpCas9n(BB)-2A-GFP | TGATGAGCACAAGGAATTCA | GAAACACAGCACACACAGAG |
| IRS1 Pro4Thr | pSpCas9n(BB)-2A-GFP | GAGGGCTCGCCATGCTGCTG | CTTCTCAGACGTGCGCAAGG |
| IRS1 Tyr465His | pSpCas9n(BB)-2A-GFP | CTCCTCACCCCTGGCTGGTG | CTGAGCAATTATATCTGCAT |
| PHLPP1 Pro298Leu | pSpCas9n(BB)-2A-GFP | GCCGATCTGCCGCTGCCCGG | GAGGCGCGCGCGCAGGGCCC |
| PHLPP1 Leu843Pro | pSpCas9n(BB)-2A-GFP | CACCAAGCTTATTGTCTCGC | GACCTGCGAGACAATAAGCT |
| PHLPP1 Glu1149Gly | pSpCas9n(BB)-2A-GFP | AGTGAGTATTGCAAAGCATG | TTGTGGTCCAGGGCAAGGCG |

**Table S2 | Overview of ssODNs used for CRISPR of mESCs.**

| Target mutation | ssODN |
|---|---|
| DEPTOR Cys102Trp | ATTTAATCCTCTGTCTTGTGTGAGACTTCTCACTCT*A*CTCTCTGTG*C*GTGCTGTGTTTCAGTGTG**G**GA*C*GAGCACAAGGAATT*T*AAGGATGTAAAACTCTTCTACCGCTTTAGGAAGGAT |
| IRS1 Pro4Thr | TATGCATACTCTTGGGCTTGCGCAGGTAGCC*G*ACCTTGCGCACGTCTGAGAAGCCATCGGTATCCGGAG**T**GCTCGCCATGCTGCTGCG*T*AGCAGAGGGAGGTGTTGAAAAACTGGGTGAG |
| IRS1 Tyr465His | GGAGCAGCCAAGGTGGAGGCTCCCTTGCCACCCATGCAGATAT**G**ATTG*GA*CAGCTCCTCCTCACCCCTGGCTGGTGG*T*GTGTGCCCCAGGGAATCTGGGGTGACACTGCGGAAGGAACTT |
| PHLPP1 Pro298Leu | CGTGGAGCCACCGCCCTCGAGCGGCACTGTTGGTGCTGT*T*CGGGGCCCTGCGCGCGCGC**T**TCCCGC*A*GATCTGCCGCTGCC*T*GGGGGCGCCTGGACGCGCTGTGCACCCCGCATCAGCCC |
| PHLPP1 Leu843Pro | GCTTATGGCAGATGAGGTGGACTTTGTGCAACATGTCACTCAGCTTGACCTGCG*T*GA*T*AATAAGC**C**TGGTGATCTAGATGCTATGATCTTCAACAACATAGAAGTTCTGCACTGTGAGAG |
| PHLPP1 Glu1149Gly | GCAGGAGCTAGACCTGACTGGAAATCCACGC*TTG*GCCCTGGACCACAAAAGCCTGG**G**GCTGCTCAAGTGAGTATTGCAAAGCATGTG*A*TCCCTCATGTGTTCACTTGCAGTGGGGTTTAC |

The target mutations are highlighted in bold, silent PAM, silent gRNA, and restriction site mutations are highlighted in italics.

**Table S3 | Overview of genotyping primer used for CRISPR of mESCS.**

| Target mutation | Forward Primer | Reverse Primer |
|---|---|---|
| DEPTOR Cys102Trp | GAAAGCTTCTCAGAGCAGTCC | CAAAATAAATAAGTAACCAAGCCAAA |
| IRS1 Pro4Thr | GTCATGGTGGGCCTTTGC | GGATCAGGCTATCTTCCTTGG |
| IRS1 Tyr465His | CTCCACGACGTTCCAAGTG | CCATGGCTCCACTTCAGACT |
| PHLPP1 Pro298Leu | GGAGCCTGGACAGGAAGAC | CCGGTCTGACACGCTCTC |
| PHLPP1 Leu843Pro | TCTGTACTTTCTCAACCTTGTTTTT | GAGATGAGGAAAGATGAGTGTCAG |
| PHLPP1 Glu1149Gly | GGCATTATAACCCTGAAGATCC | GCATCACAGTCATGTGACCA |

**Table S4 | Overview of primary antibodies used for western blotting on mESCS.**

| Antigen | Source | Dilution | Vendor #ID |
|---|---|---|---|
| Phospho-AKT (Thr308) | Rabbit | 1:1,000 | Cell Signaling #9275 |
| Phospho-AKT (Ser473) | Rabbit | 1:1,000 | Cell Signaling #4060 |
| Total AKT | Rabbit | 1:1,000 | Cell Signaling #9272 |
| Phospho-p70 S6K (Thr389) | Rabbit | 1:1,000 | Cell Signaling #9205 |
| p70 S6K total | Rabbit | 1:1,000 | Cell Signaling #9202 |
| Phospho-p44/42 MAPK (Erk1/2) (Thr202+Tyr204) | Rabbit | 1:1,000 | Cell Signaling #4370 |
| p44/42 MAPK (Erk1/2) total | Rabbit | 1:1,000 | Cell Signaling #4695 |
| Phospho-MEK1/2 (Ser217/221) | Rabbit | 1:1,000 | Cell Signaling #9154 |
| MEK1/2 total | Rabbit | 1:1,000 | Cell Signaling #9122 |
| GAPDH | Rabbit | 1:1,000 | Cell Signalling #2118 |

Antibodies were ordered from Cell Signaling Technology.

**Table S5 | Overview of TaqMan probes used for qPCRs on mESCs.**

| Gene | TaqMan Assay ID |
|---|---|
| *Ets1* | Mm01175819_m1 |
| *Ets2* | Mm00468973_m1 |
| *Etv6* | Mm01261325_m1 |
| *Foxo3* | Mm01185722_m1 |
| *c-Myc* | Mm00487804_m1 |
| *Gapdh* | Mm99999915_g1 |
| *Cox1* | Mm04225243_g1 |
| *Rnr2* | Mm04260181_m1 |
| *18S* | Hs9999999901_s1 |
